# Translation factor mRNA granules direct protein synthetic capacity to regions of polarized growth

**DOI:** 10.1101/447680

**Authors:** Mariavittoria Pizzinga, Christian Bates, Jennifer Lui, Gabriella Forte, Fabián Morales-Polanco, Emma Linney, Barbora Knotkova, Beverley Wilson, Clara A. Solari, Luke E. Berchowitz, Paula Portela, Mark P. Ashe

## Abstract

mRNA localization serves key functions in localized protein production making it critical that the translation machinery itself is present at these locations. Here we show that translation factor mRNAs are localized to distinct granules within yeast cells. In contrast to many mRNP granules, such as P-bodies and stress granules, which contain translationally repressed mRNAs, these granules harbor translated mRNAs under active growth conditions. The granules require Pab1p for their integrity and are inherited by developing daughter cells in a She2p/ She3p dependent manner. These results point to a model where roughly half the mRNA for certain translation factors are specifically directed in granules toward the tip of the developing daughter cell where protein synthesis is most heavily required, which has particular implications for filamentous forms of growth. Such a feedforward mechanism would ensure adequate provision of the translation machinery where it is to be needed most over the coming growth cycle.

**Summary:** This study shows that mRNAs encoding a range of translation factors are localized to granules that get transported into the yeast daughter cell using the She2p/She3p machinery. This likely supports an intensification of protein synthetic activity to facilitate apical polarized growth.

## Introduction

mRNA localization serves to regulate spatiotemporal protein production playing critical functions across physiology. These functions include cell and tissue differentiation, cellular polarization, and protein targeting to organelles and membranes (Holt and Bullock, 2009; Pizzinga and Ashe, 2014). mRNAs encoding key determinants in developing oocytes/ embryos were among the first examples of localized mRNAs to be discovered. Prominent examples include: *β-actin* mRNA in Ascidian embryos (Jeffery et al., 1983), *Vg1* in *Xenopus* oocytes (Melton, 1987) and *bicoid* mRNA in the *Drosphila* oocyte (Berleth et al., 1988). Further cases were described in neuronal cells where specific mRNAs were found in dendritic and/or axonal regions providing neurons with the flexibility of structure and function required for synaptic plasticity (Garner et al., 1988; Miyashiro et al., 1994). Even in unicellular eukaryotes like yeast, mRNAs such as *ASH1* have been found to deliver a polarity of phenotype between the mother and daughter cell (Long et al., 1997; Takizawa et al., 1997).

More recent assessments of the role of mRNA localization in the control of gene expression suggest that, rather than being restricted to just a handful of mRNAs, regulated localization is remarkably widespread. For instance, studies in the *Drosophila* embryo have shown that approximately 70% of expressed mRNAs are localized in some manner (Lecuyer et al., 2007). Furthermore, large numbers of localized mRNAs have also been observed in the *Drosophila* ovary (Jambor et al., 2015), neuronal axon growth cones (Zivraj et al., 2010) and dendrites (Cajigas et al., 2012). Even in yeast, the localization of mRNA appears much more commonplace than previously anticipated: mRNAs encoding peroxisomal, mitochondrial and ER proteins, as well as mRNAs for general cytoplasmic proteins are localized (Fundakowski et al., 2012; Gadir et al., 2011; Lui et al., 2014; Schmid et al., 2006; Zipor et al., 2009). However, studies at the mRNA-specific level have uncovered a number of key principles that have been found to resonate across many mRNA localization systems in various eukaryotic species.

The ‘prototype’ mRNA in yeast studies was the *ASH1* mRNA, which localizes to the tip of the daughter cell in order to repress yeast mating type switching, such that this process only occurs in mother cells (Singer-Kruger and Jansen, 2014). *ASH1* mRNA localization was found to rely upon actin cables and a specific myosin, Myo4p. In addition, the RNA-binding protein She2p interacts with the mRNA and, via the She3p scaffold protein, targets the mRNA to Myo4p (Singer-Kruger and Jansen, 2014). More generally, cytoskeletal microtubules or actin filaments, as well as appropriate motor proteins, have been identified as a common feature of many mRNA localization mechanisms (Lopez de Heredia and Jansen, 2004).

The *ASH1* mRNA localization system and its role in mating type switching highlights another key feature of many mRNA localization events. That is, since any inappropriate expression of Ash1p essentially compromises the difference between the mother and daughter cell, the system is wholly reliant upon the translational repression of *ASH1* mRNA during transit. Similar tight regulation of protein synthesis during transit has been identified for mRNAs in other systems such as morphogenetic gradient formation in *Drosophila* oocytes/ embryos (Lasko, 2012; St Johnston, 2005). Equally such regulatory processes rely upon specific derepression of the mRNA, once it reaches its final destination (Besse and Ephrussi, 2008). In the case of *ASH1* mRNA, two mechanisms of translational derepression have been proposed involving Puf6p and Khd1p, respectively (Deng et al., 2008; Paquin et al., 2007).

As well as at the mRNA-specific level, translation repression can also occur at a more global level: for instance as a response to stress (Simpson and Ashe, 2012; Spriggs et al., 2010). Such widespread repression also has defined consequences in terms of mRNA localization: translationally repressed mRNA can be transferred to mRNA processing bodies (P-bodies) or stress granules (Brengues et al., 2005; Hoyle et al., 2007; Kedersha et al., 2000; Mollet et al., 2008; Simpson et al., 2014). P-bodies house many mRNA degradation components and have been considered as sites of mRNA decay (Jain and Parker, 2013), although a more recent evaluation of P-body function in human cells has favored a more dominant role in mRNA storage (Hubstenberger et al., 2017). Stress granules harbor a variety of RNA-binding proteins/ translation factors and are thought of as sites of mRNA storage or triage (Anderson and Kedersha, 2008; Buchan and Parker, 2009). Recent studies have highlighted that these bodies adopt a more dynamic liquid structure than previously appreciated, such that enzymatic activities and protein refolding might be conceivable within the body (Aguzzi and Altmeyer, 2016; Mitrea and Kriwacki, 2016; Sfakianos et al., 2016). Critically, both P-bodies and stress granules can be induced by cellular stresses that bring about the robust repression of translation initiation (Hoyle et al., 2007; Kedersha et al., 2005; Teixeira et al., 2005; Wilczynska et al., 2005).

Recently, we have found that mRNAs can localize to granules even in rapidly growing cells (Lui et al., 2014). It appears that, at least for granules harboring mRNAs encoding components of the glycolytic pathway, active translation of mRNAs is occurring at these sites. This is suggestive that the mRNA granules might represent the kind of liquid, dynamic structure described above. Intriguingly, these mRNA granules also appear to seed the formation of P-bodies after stress and might represent sites where the fate of similar classes of mRNA is co-ordinated (Lui et al., 2014).

In this current paper, we have investigated the localization of another class of mRNA in actively growing cells. We find that mRNAs encoding translation factors localize to mRNA granules that are different to those previously described to carry glycolytic mRNAs; they are fewer in number and display distinct inheritance patterns. Indeed, translation factor mRNA granules are specifically inherited by daughter cells and appear to play a role in focusing translational activity at sites of polarized growth during yeast filamentous growth. Overall, the protein synthetic capacity of a cell accumulates at specific sites via the localization of key mRNAs to facilitate polarized growth.

## Results

### Translation factor mRNAs are localized in actively growing yeast

Previous work from our laboratory has identified yeast mRNAs that are localized to and translated in granules during active cell growth (Lui et al., 2014). More specifically, two glycolytic mRNAs, *PDC1* and *EN02,* were investigated using MS2-tagging of endogenous mRNAs (the m-TAG system) and Fluorescent *in situ* hybridization (FISH) to reveal localization to such granules. In actively growing cells, these mRNAs co-localized to 10-20 granules per cell, whereas following a switch to translation repression conditions, the granules were observed to coalesce then recruit P-body components (Lui et al., 2014). In these studies, the *TIF1* mRNA, which encodes the translation initiation factor elF4Al, was also identified as localized in unstressed cells. However, in this case, fewer granules per cell (<5) were observed (Lui et al., 2014).

In order to expand our understanding of the localization of mRNAs encoding the protein synthetic machinery, a range of mRNAs encoding translation factors were selected and tagged using the m-TAG system. mRNAs were selected that produce proteins with a range of abundances within cells and that participate across the three phases of translation: initiation, elongation and termination (Fig. IB). The m-TAG technique involves the precise addition of *MS2* stem loops into the 3’ UTR of genes at their genomic loci, then coexpression of GFP fused to the MS2 coat protein (Haim-Vilmovsky and Gerst, 2009). Similar MS2-based GFP tethering systems have been widely used in yeast and other cells to study many aspects of RNA biology. The key advantage of this technology is that it allows the localization of mRNA to be studied in live cells (Buxbaum et al., 2015).

Intriguingly, all the investigated mRNAs that encode components of the translation machinery localized to granules in unstressed cells (Fig. 1A). Critically, under these highly active growth conditions, P-bodies and stress granules are not evident (Lui et al., 2014)(Fig. 3D). Hence, the mRNA localization observed is not related to P-bodies and stress granules that form under defined stress conditions in yeast (Buchan et al., 2011; Grousl et al., 2009; Hoyle et al., 2007; Iwaki and Izawa, 2012; Shah et al., 2016; Teixeira et al., 2005). It is important to note that the observed mRNA localization patterns for the translation factor mRNAs do not necessarily represent the norm: many other mRNAs have a broad cytoplasmic localization profile (Lui et al., 2014). For instance, *NPC2* is used as a control here to illustrate this point (Fig. 1A).

**Figure 1.**
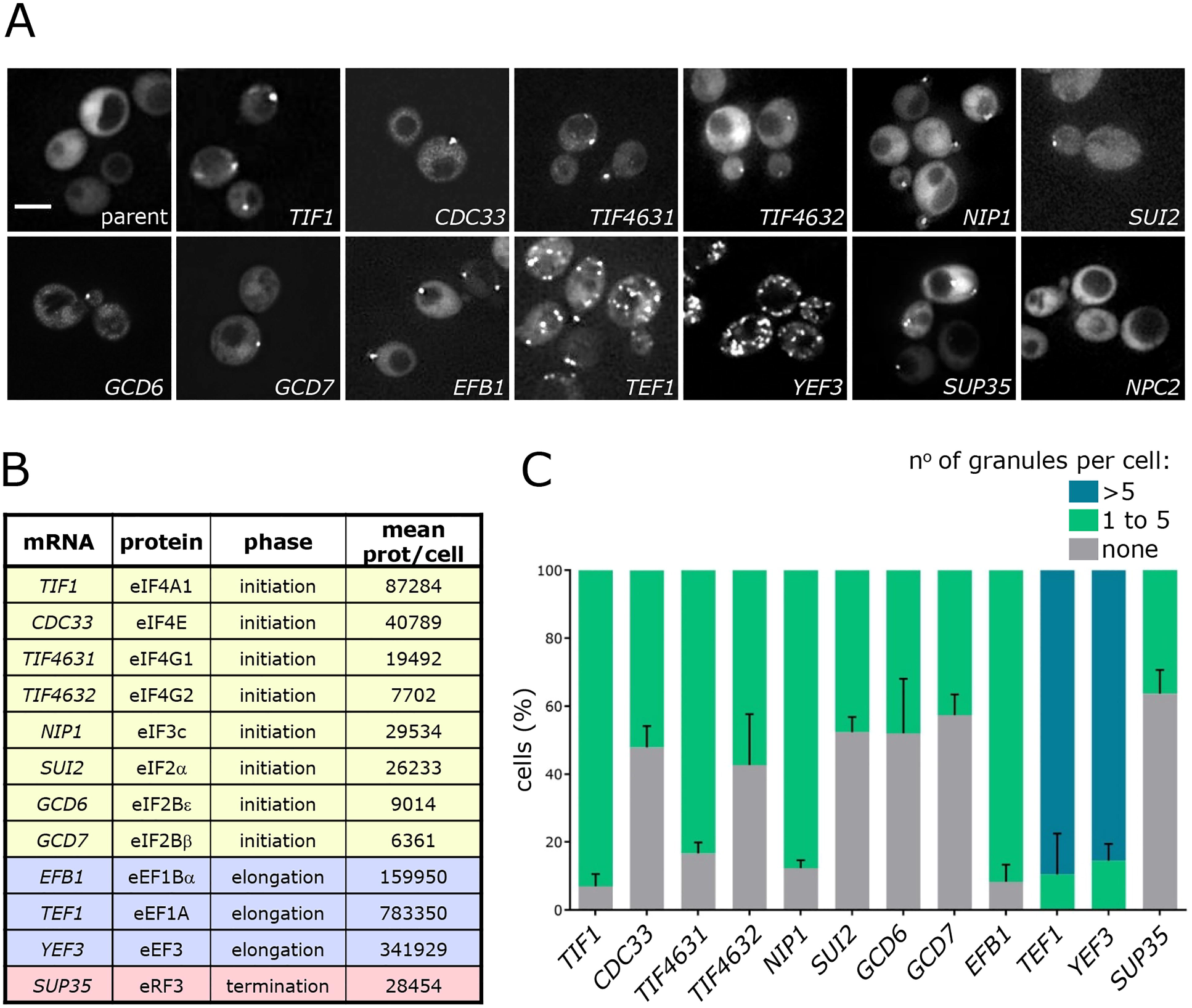
Translation factor mRNAs localize to cytoplasmic granules in exponentially growing *S. cerevisiae.* (A) Z-stacked images of strains expressing specific *MS2-*tagged mRNAs as labelled and the MS2 coat protein GFP fusion. Scale bar = 4 μm. (B) List of the tagged mRNAs, the proteins they encode, the translation phase they are involved in and the mean number of proteins per cell (taken from Ho et al., 2017). (C) Chart showing the percentage of cells with 1-5, over 6 or no granules per cell. Error bars indicate +SD over three biological replicates.

**Figure 3.**
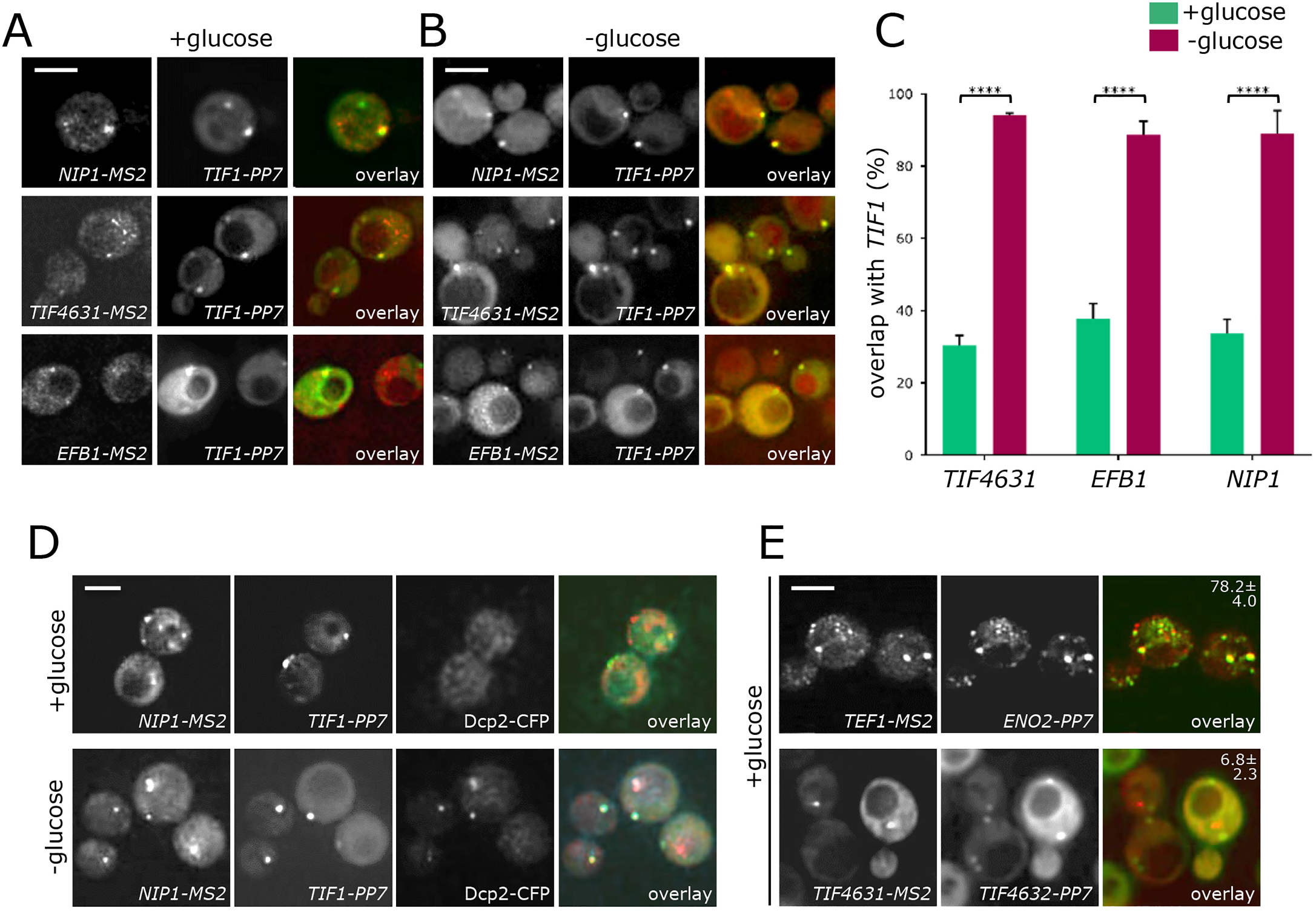
Co-localization analysis of *MS2-* and *PP7*-tagged strains. Z-stacked images showing localization of *NIP1-MS2, TIF4631-MS2* and *EFB1-MS2* (via the co-expressed MS2 coat protein mCherry fusion) relative to *TIF1-PP7* (visualised using co-expressed PP7 coat protein GFP fusion). (A) Actively growing in SCD media and (B) following glucose starvation for 10 min. (C) Chart showing the percentage of observable *NIP1-MS2, EFB1-MS2* or *TIF4631-MS2* granules co-localising with *TIF1-PP7* granules in SCD media (green) and following glucose starvation for 10 min (magenta). **** p<0.0001. Error bars are indicate +SD. (D) Z-stacked images of *NIP1-MS2* and *TIF1-PP7* mRNAs relative to a P-body marker Dcp2p tagged with CFP both in SCD media and after glucose depletion for 10 min (E) Z-stacked images of mRNAs: *TEF1-MS2* relative to *EN02-PP7* and *TIF4631-MS2* relative to *TIF4632-PP7.* The percentage of observable MS2-tagged mRNA co-localizing with the PP7-tagged mRNA is indicated ±SD. Scale bars = 4 μm.

Even though all the translation factor mRNAs are localized, variation in the pattern of localization in terms of the number of granules per cell is evident (Fig. 1A and C). Most of the mRNAs, including those encoding all the translation initiation factors, the eRF3 *(SUP35)* translation termination factor and the eEFlBa *(EFB1)* elongation factor localize to less than 5 granules per cell (Fig. 1A and C). In contrast, the two other tested elongation factor mRNAs, eEF1A *(TEF1)* and eEF3 *(YEF3),* localize to many more granules per cell: in the order of 10-20. The higher number of mRNA granules is more similar to that previously observed for two yeast glycolytic mRNAs (Lui et al., 2014). When expression profiles were evaluated using the SPELL algorithm (version 2.0.3r71) (Hibbs et al., 2007), which compares expression profiles to identify similarly regulated genes across a plethora of transcriptomic experiments, the translation elongation factor genes were identified as more similar to glycolytic genes than to genes encoding the rest of the translation machinery. It therefore seems that the expression of these translation elongation factor mRNAs is co-regulated with mRNAs of the glycolytic pathway.

It is also noticeable, when the levels of the various mRNAs were assessed by quantitative reverse transcriptase PCR, that the *TEF1* (eEF1A) and *YEF3* (eEF3) mRNAs are the most abundant (Fig. SI). This highlights the possibility that mRNA abundance may play a role in the propensity of an mRNA to enter granules. However, the abundance measurements for other mRNAs do not equate with their presence in granules. For instance, for the other translation factor mRNAs in granules, abundance can vary from relatively low: *TIF4631* (elF4Gl) and *TIF4632* (eiF4G2) mRNAs, to *TIF1* (elF4Al) mRNA, which is nearly as abundant as the translation elongation factor mRNAs (Fig. S1). Even though there is nearly a 100-fold difference between these extremes, the localization of the mRNAs is remarkably similar. This is suggestive that the presence of an mRNA within RNA granules or the pattern of those mRNA granules is not merely reflective of the overall abundance of an mRNA.

Recent studies have highlighted the potential impact of the addition of *MS2* stem loops to aspects of mRNA fate (Garcia and Parker, 2015; Garcia and Parker, 2016; Haimovich et al., 2016; Heinrich et al., 2017). Indeed, in our own previous studies, *MS2* stem loops were found to decrease the levels of both the *MFA2* mRNA and the *PGK1* mRNA (Lui et al., 2014; Simpson et al., 2014). In this current study, we have assessed the impact of *MS2* stem loop insertion and expression of the MS2-GFP fusion protein in the m-TAG strains relative to the untagged mRNA in the parent strain (Fig. SI). This analysis suggests that the *MS2* stem loops and fusion protein can have a complex and variable impact upon the production and stability of an mRNA. For some mRNAs, such as *CDC33* (elF4E), *EFB1* (eEF1Bα), *GCD6* (elF2Bε) and *GCD7* (elF2Bβ) the introduction of the MS2 system leads to a significant decrease in mRNA levels. For others, such as *SUI2* (elF2α), *TIF4631,* (elF4Gl), and *TIF4632* (elF4G2), the MS2 system has little significant effect on the overall level of the mRNA. In addition, there are a number of mRNAs where the MS2 system has an intermediate effect. It is unclear why the introduction of this system should reduce mRNA levels and it is possible that multiple factors are at play. For instance, it is plausible that the introduction of the stem loops or just generally alterations within the 3’ UTR would impact upon the production and 3’ end processing of the mRNA, or it could alter mRNA stability. This is not particularly surprising given that one well-established strategy for reducing essential gene function in yeast is to insert a marker into the 3’ UTR of a gene of interest (Breslow et al., 2008).

Given the variability in the effects caused by introduction of the MS2 system to an mRNA and concerns regarding the integrity of mRNAs that are bound by the MS2-GFP fusion protein, it was important to assess mRNA localization using another independent technique such as fluorescent *in situ* hybridization (FISH). In previous studies, we have used FISH to show that the m-TAG system can reflect the genuine localization of endogenous mRNAs in yeast (Lui et al., 2014). Here, we adapted a single molecule FISH technique (Tsanov et al., 2016) for use in yeast to generate a high resolution profile of the location of endogenous translation factor mRNAs (Fig. 2A). smFISH is more sensitive than m-TAG in that both large multi-mRNA granules and smaller single mRNA foci are observed (Fig. 2A). For all the mRNAs examined using smFISH, the number of large multi-mRNA granules per cell correlates well with the numbers of granules per cell obtained using the MS2 system (Fig. 2A). Even for *YEF3* (eEF3) where many mRNA granules were observed with the MS2 system, numerous large mRNA granules were also observed with smFISH. From the smFISH data, it is possible to estimate the average number of mRNA molecules for an individual mRNA species per cell.

**Figure 2.**
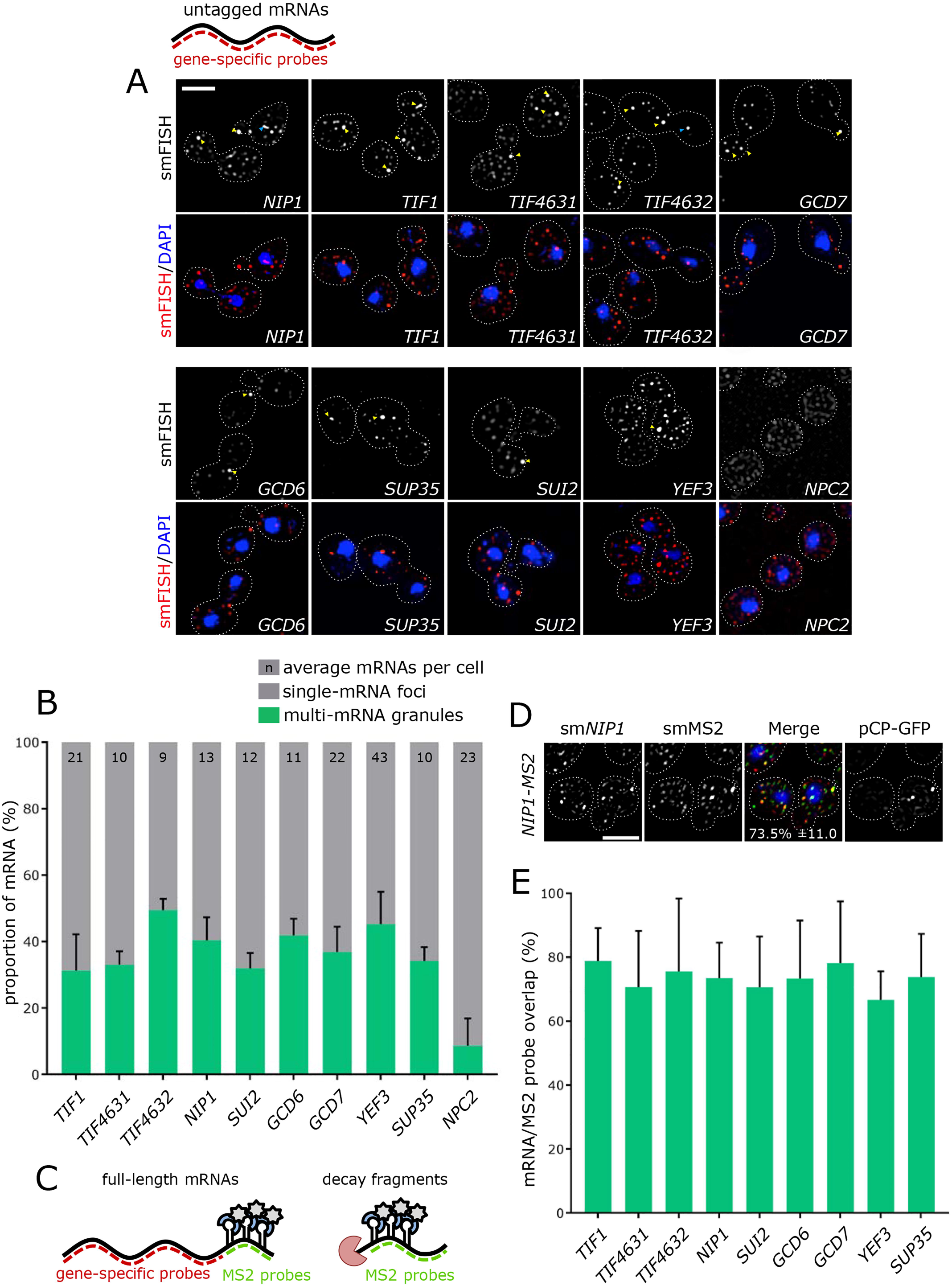
smFISH recapitulates mRNA localization observed with the m-TAG system. (A) Z-stacked images of smFISH performed for the genes indicated, using a W303-1A wild type strain expressing endogenous mRNAs. (B) Bar chart showing the proportion of mRNA in either single mRNA foci (<2.5 mRNAs per spot), or granule foci (>2.5 mRNAs per spot). Numbers indicate the average number of mRNA molecules observed per cell based on both single mRNA and granule foci. (C) Diagram depicting the experimental approach. By using a combination of specifically labelled probes that anneal to the gene body and *MS2* region, it is possible to observe whether pCP-GFP signal arises from full-length mRNAs (left), or the aggregation of 3’ decay fragments (right). (D) Example Z-stacked image of the strain expressing MS2-tagged *NIP1,* visualising the gene body (smNIPl), *MS2* loops (smMS2) and pCP-GFP signal. (E) Bar chart quantifying the degree of overlap between MS2-GFP foci and gene body foci for a range of MS2-tagged mRNAs, as indicated. Error bars indicate +SD. Scale bars = 3 μm.

Such estimates compare very favourably with the number of mRNA molecules per cell that have been calculated from two RNA-seq based studies (Fig. S2A and B) (Lahtvee et al., 2017; Lawless et al., 2016). This analysis also revealed the number of molecules of an mRNA species that are present in the large granules as a proportion of total (Fig. 2B). As a result of this analysis, we conclude that roughly half of the translation factor mRNAs in each cell are present in large granules, and the other half are present as single molecules. Interestingly for the *NPC2* mRNA, which was not observed in granules using the MS2 system, a much lower proportion of total mRNA was present in large granules using smFISH. These data show that endogenous translation factor mRNAs localize to large cytoplasmic granules and that the number of large granules is similar to that observed in the live cell m-TAG system.

In order to further explore the relationship between the profiles observed for the m-TAG and smFISH techniques, smFISH was performed in the m-TAG strains comparing the localization profile observed for probes against the body of the mRNA versus probes to the *MS2* stem loop portion (Fig. 2C and D). This comparison reveals a high degree of overlap with greater than 75% of signal observed with the *MS2* probe overlapping with signal for the mRNA body probe (Fig. 2D and E). Furthermore, significant overlap was also observed with the GFP signal generated from the MS2-GFP fusion that is expressed in these yeast strains (Fig. 2D) and that has been observed in live cells (Fig. 1). However, what is clear from this analysis is that only the most intense foci from the smFISH contain discernible GFP signal from the MS2-GFP fusion (Fig. S2D). These data support the interpretation that the MS2 system does not detect single molecule mRNAs and only reveals larger multi-mRNA granules, as the MS2-GFP fusion is only detected for granules which have higher smFISH signal intensity (Fig S2C and D). However, the key point of this experiment is that where signal from the MS2-GFP was identified, signal from the mRNA body was also evident (>90%). Therefore, it appears that the MS2 system can faithfully reproduce endogenous mRNA localization patterns and can report the presence of full-length mRNAs but in these experiments not at single molecule resolution. In summary, these results further support the hypothesis that mRNA localization plays an important role in determining the fate of mRNAs encoding components of the translation machinery.

### mRNA granules harbor a complex mix of translation factor mRNAs

A key question relating to the mRNA granules described above is whether each granule within cells contains mRNAs for most translation factors, or whether numerous RNA granules exist with a more variable mRNA composition. To address this question in live cells, we used a scheme to allow the localization of different mRNAs to be cross-compared. This scheme combines the *MS2* mRNA localization system with an analogous yet discrete system in terms of specificity: the *PP7* mRNA localization system (Hocine et al., 2013; Lui et al., 2014). Therefore, strains were generated with *PP7*-tagged *TIF1* (elF4Al) mRNA and *MS2* stem loops in the 3’ UTR of another translation factor mRNA. Two fusion proteins were coexpressed: PP7 coat protein fused to GFP and the MS2 coat protein fused to mCherry. This allowed the simultaneous assessment of two different mRNAs within the same live cell (Fig.3A).

A comparison of the degree of overlap for the observed granules revealed that for each of the mRNAs *TIF4631* (elF4G1), *NIP1* (elF3c) and *EFB1* (eEF1Bα), approximately 30% of mRNA granules also contained the *TIF1* (elF4A1) mRNA (Fig. 3C). Control experiments reveal that this degree of co-localization is not due to crosstalk between the fluorescent channels (Fig. S3). We consider this overlap highly significant, as in previous studies where we have assessed the overlap between a glycolytic mRNA *(PDC1)* and a translation factor mRNA *(TIF1)* we found no overlap (Lui et al., 2014). Moreover, comparison of the localization of *TIF4631* (elF4Gl) mRNA with the *TIF4632* (elF4G2) mRNA also exhibited low levels of colocalization (Fig. 3E). As well as highlighting the significance of the degree of overlap observed for other combinations (Fig. 3A), this result indicates that not every mRNA is colocalized to the same set of granules.

In contrast to the mRNAs studied above, which do not co-localize with glycolytic mRNAs (Lui et al., 2014), there seems to be a significant degree of co-localization between the elongation factor-encoding mRNAs *(TEF1* and *YEF3),* and the glycolytic mRNA *EN02* (Fig. 3E). Previously *EN02* was shown to overlap almost perfectly with PDC1-containing granules and it was suggested that these granules might represent a site where mRNAs of similar function are co-regulated (Lui et al., 2014). The fact that translation elongation factor mRNAs are also localized to the same granules further correlates with the transcriptional co-regulation, mentioned above, between these translation elongation factor mRNAs and mRNAs encoding glycolytic components that is evident from analysis using the SPELL algorithm (version 2.0.3r71) (Hibbs et al., 2007).

In order to corroborate these live cell co-localization studies, dual mRNA smFISH experiments were undertaken to investigate the degree of co-localization for various endogenous mRNAs (Fig. S4). As in the MS2/PP7 live cell experiments, the degree of colocalization for *TIF1* versus *NIP1* and *TIF1* versus *TIF4631* was in the range 30-40% (Fig. S4). In contrast, much lower co-localization was observed when *TIF4631* and *TIF4632* mRNAs were compared (Fig. S4). These smFISH results on endogenous mRNAs in fixed cells almost precisely parallel the observation made using the MS2/ PP7 system in live cells.

Therefore, it is clear that not every translation factor mRNA is contained in every granule; for instance, *TIF4631* and *TIF4632* mRNAs appear almost mutually exclusive. Instead, the results above support a model where a complex cocktail of translation factor mRNAs are housed within numerous mRNA granules.

### Translation factor mRNA granules coalesce to form P-bodies after stress

Previous work has suggested that mRNA granules carrying the *PDC1* and *EN02* glycolytic mRNAs coalesce to seed the formation of P-bodies under glucose starvation conditions (Lui et al., 2014). To address the fate of the granules carrying translation factor mRNAs during P-body formation, the PP7/MS2 co-localization strains were again utilized under the rapid glucose depletion conditions that are known to induce P-bodies (Fig. 3B). In this case, after 10 minutes of glucose depletion approximately 90% of granules contained both *TIF1* mRNA and the relevant MS2-tagged mRNA *(TIF4631, NIP1* or *EFB1)* (Fig. 3C). These data are consistent with a view that the translation factor mRNA granules also coalesce during the formation of P-bodies.

However, in order to directly assess whether these coalesced RNA granules are in fact P-bodies, the *NIP1* and *TIF1* mRNAs were evaluated at the same time as a CFP-tagged P-body marker protein Dcp2p (Fig. 3D). Consistently with previous observations (Lui et al., 2014), the P-body marker Dcp2p localizes broadly throughout the cytosol and does not overlap with the RNA granules in actively growing cells (Fig 3D). However, 10 minutes after glucose starvation both the *TIF1* and *NIP1* mRNAs as well as Dcp2p are found in the same granules (Fig. 3D). These experiments collectively support a view where the translation factor mRNA granules contribute to the formation of P-bodies in a similar manner to the RNA granules carrying glycolytic mRNAs (Lui et al., 2014).

### mRNA translation is a requirement for translation factor mRNA localization to granules

Previous work has suggested that mRNA granules can serve as sites of mRNA translation in actively growing yeast (Lui et al., 2014). To investigate whether translation of a specific mRNA affects its capacity to localize to granules, a well-characterised stem loop (ΔG value of −41 kcal/mol) was inserted into the 5’ UTR of the *NIP1* mRNA. This stem loop has previously been widely used to reduce translation of specific mRNAs by limiting scanning of the 43S preinitiation complex through to the AUG START codon without impacting upon the stability of the mRNA (Palam et al., 2011; Vattem and Wek, 2004). In this case, the MS2-tagged *NIP1* mRNA was derived from a plasmid rather than the genome within the yeast strain. A direct comparison of *NIP1* mRNA localization from the plasmid versus genomic system revealed little difference in the localization to granules or number of granules per cell (Fig. S5). The insertion of a stem loop into the *NIP1* 5’UTR significantly reduced the capacity of the *NIP1* mRNA to enter the RNA granules (Fig. 4A and D). Critically, the insertion of the stem loop did not significantly alter the level of the *NIP1* mRNA expressed from these plasmid constructs (0.165 ±0.034 for *NIP1-MS2* versus 0.161 ±0.022 for *sl-NIP1-MS2* relative to *ACT1* mRNA). These data suggest that translation of the mRNA might be important for its localization.

**Figure 4.**
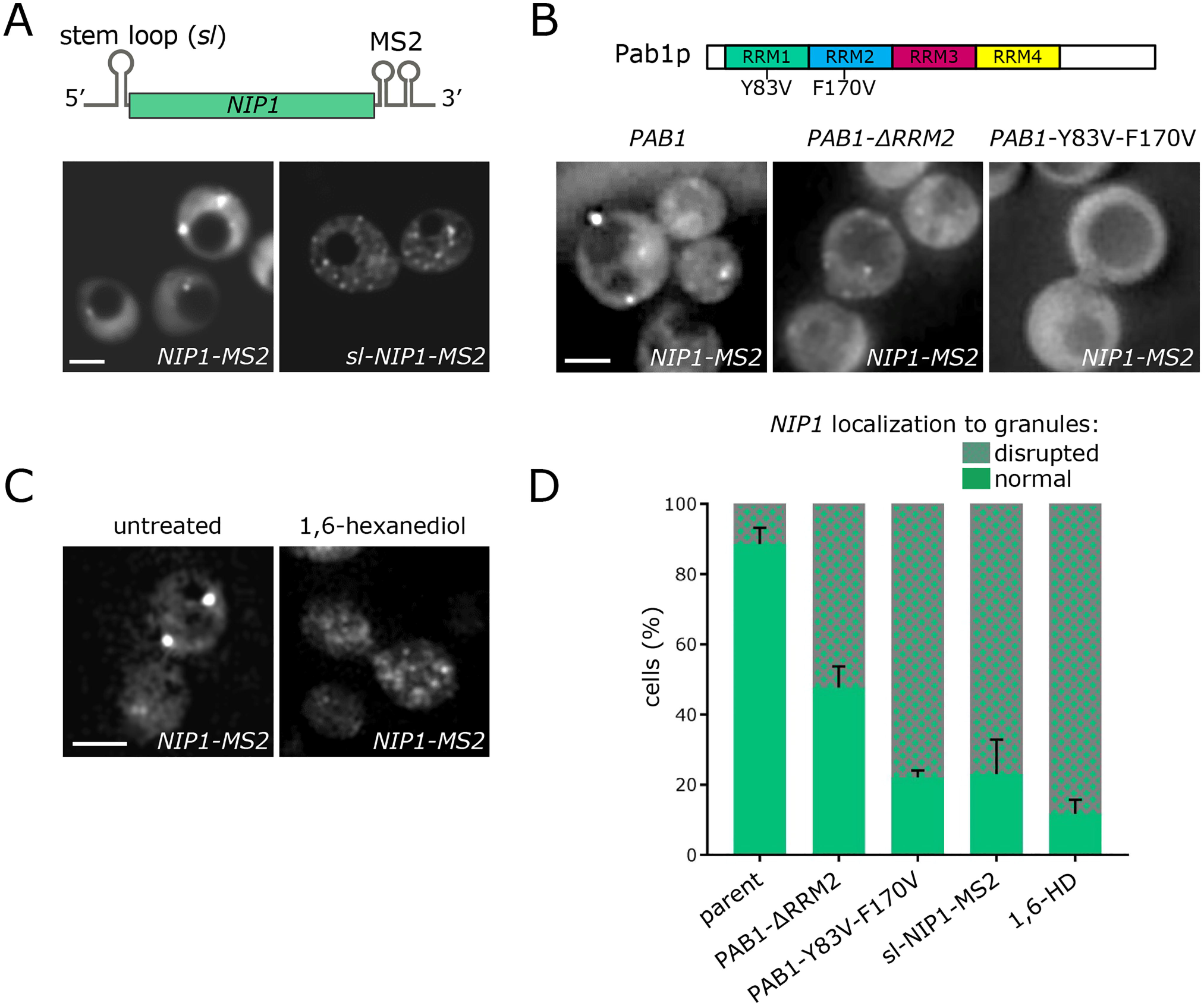
Translation is required for correct localization of translation factor mRNAs to granules. (A) A schematic of *NIP1-MS2* construct and the inserted 5’ UTR stem loop to limit translation initiation. Z-stacked images are shown below for *NIP1* mRNA localization from constructs either with *(sl-NIPl-MS2)* or without *(NIP1-MS2)* the stem loop. Scale bar = 5 μm.(B) A schematic of Pablp and the position of the two point mutations, which impact upon translation initiation. Z-stacked images are shown below for *NIP1-MS2* mRNA localization in *pab1Δ* strains bearing various *PAB1* plasmids: wild type *PAB1, PAB1* lacking the RRM2 region or *PAB1* carrying the Y83V and F170V point mutations. Scale bar = 4 μm. (C) Z-stacked images of *NIP1-MS2* mRNA in either untreated cells or after treatment with 10% 1,6-hexanediol for 30 min. Scale bar = 4 pm. (D). Bar chart depicting quantitation of the impact of stem loop insertion, *PAB1* mutation or hexanediol treatment on the integrity of the *NIP1* mRNA granules. Error bars indicate +SD.

Further evidence that translation might be important for the localization of the mRNAs comes from investigations of specific mutations in the poly(A) binding protein, Pablp. Pablp is an RNA-binding protein with a characteristic set of four RNA recognition motifs and a C-terminal domain (Kessler and Sachs, 1998). Pab1p interacts with the mRNA poly(A) tail and elevates rates of translation initiation (Sachs et al., 1997). One mechanism by which Pab1p achieves this is via promotion of a ‘closed loop complex’ via contact with the translation initiation factor, eiF4G (Costello et al., 2015; Wells et al., 1998). The RRM2 domain of Pab1p has proved critical for binding eiF4G during the formation of the closed loop structure and thus in promoting translation initiation (Kessler and Sachs, 1998). Intriguingly, the *NIP1* mRNA, while localizing to granules in strains with wild type *PAB1,* becomes mislocalized in the *PAB1-ARRM2* yeast mutant strain (Fig. 4B and D), but not in strains in which other domains of *PAB1* were deleted (data not shown). In addition, a double point mutant was tested, which carries alterations to two key aromatic residues in RRM1 and RRM2 *(PAB1-* Y83V, F170V). This mutant Pab1p retains the capacity to bind elF4G but cannot effectively bind poly(A) or promote translation initiation (Kessler and Sachs, 1998). Once again, in this mutant, the granule specific localization of the *NIP1* mRNA is abrogated. Overall across a series of *PAB1* mutants either lacking the various domains or carrying key mutations, only those impacting upon translation affected the localization of the translation factor mRNA to granules (Fig. 4B, data not shown). Once again, these results are consistent with mRNA translation being important for localization to granules in actively growing cells.

### Translation factor mRNAs are likely translated within the granules

The majority of granule-associated mRNAs are present as a response to stress (e.g. P-bodies and stress granules) or as part of a finite control of protein expression (e.g. *ASH1* or *bicoid* mRNA localization). As such, these mRNAs enter granules in a translationally repressed state (Besse and Ephrussi, 2008). In contrast, recent work from our lab suggests that for two glycolytic mRNAs, the RNA granules observed under actively growing conditions are associated with active translation (Lui et al., 2014). The stem loop insertion and *PAB1* mutant data described above suggest that a similar scenario might exist for the translation factor mRNAs.

In order that a complex and dynamic procedure such as protein synthesis can occur in an RNA granule, the components in the granule would need to be present in a dynamic assembly, such as liquid droplets. A number of non-membrane bound compartments have recently been identified to form as a result of liquid-liquid phase separation (Aguzzi and Altmeyer, 2016). The flexible series of fluctuating weak interactions that hold together such droplets make enzymatic activity within a non-membrane bound compartment plausible, whereas it is difficult to envisage such activity within more stably aggregated granules (Mitrea and Kriwacki, 2016; Sfakianos et al., 2016). In order to gain hints as to whether the RNA granules carrying translation factor mRNAs are liquid droplets, 1,6-hexanediol was used. This reagent has been established to disrupt phase-separated liquid droplets while solid particles are unaffected (Kroschwald et al., 2015). Treatment of yeast cells with this reagent led to almost complete disruption of granules bearing the *NIP1* mRNA (Fig. 4C and D). This reagent also led to the inhibition of translation initiation (Fig. S6), as well as the disruption of other cytoskeletal functions in cells (Wheeler et al., 2016). Whether these effects occur as a result of the general disruption of processes requiring particles in the liquid state is currently unknown. Clearly, if sufficient mRNAs are translated in such particles their disruption could conceivably lead to the translation inhibitory effects that we observe.

In order to assess whether active translation of translation factor mRNAs can occur within granules, a recently described technique called ‘Translating RNA Imaging by Coat protein Knock-off or ‘TRICK’ (Halstead et al., 2015) was adapted for use in yeast. TRICK relies upon the insertion of *PP7* stem loops within an mRNA’s coding sequence, upstream of the STOP codon; and *MS2* stem loops downstream of the STOP codon within an mRNA’s 3’ UTR. If the *TRICK*-tagged mRNA is not translated, the PP7 coat protein fused to GFP and the MS2 coat protein fused to mCherry bind simultaneously. Whereas upon translation, the PP7 coat protein is displaced as ribosomes translate the coding region where the PP7 stem loops are located, resulting in the mRNA only binding the MS2-CP-mCherry (Fig. 5A).

**Figure 5.**
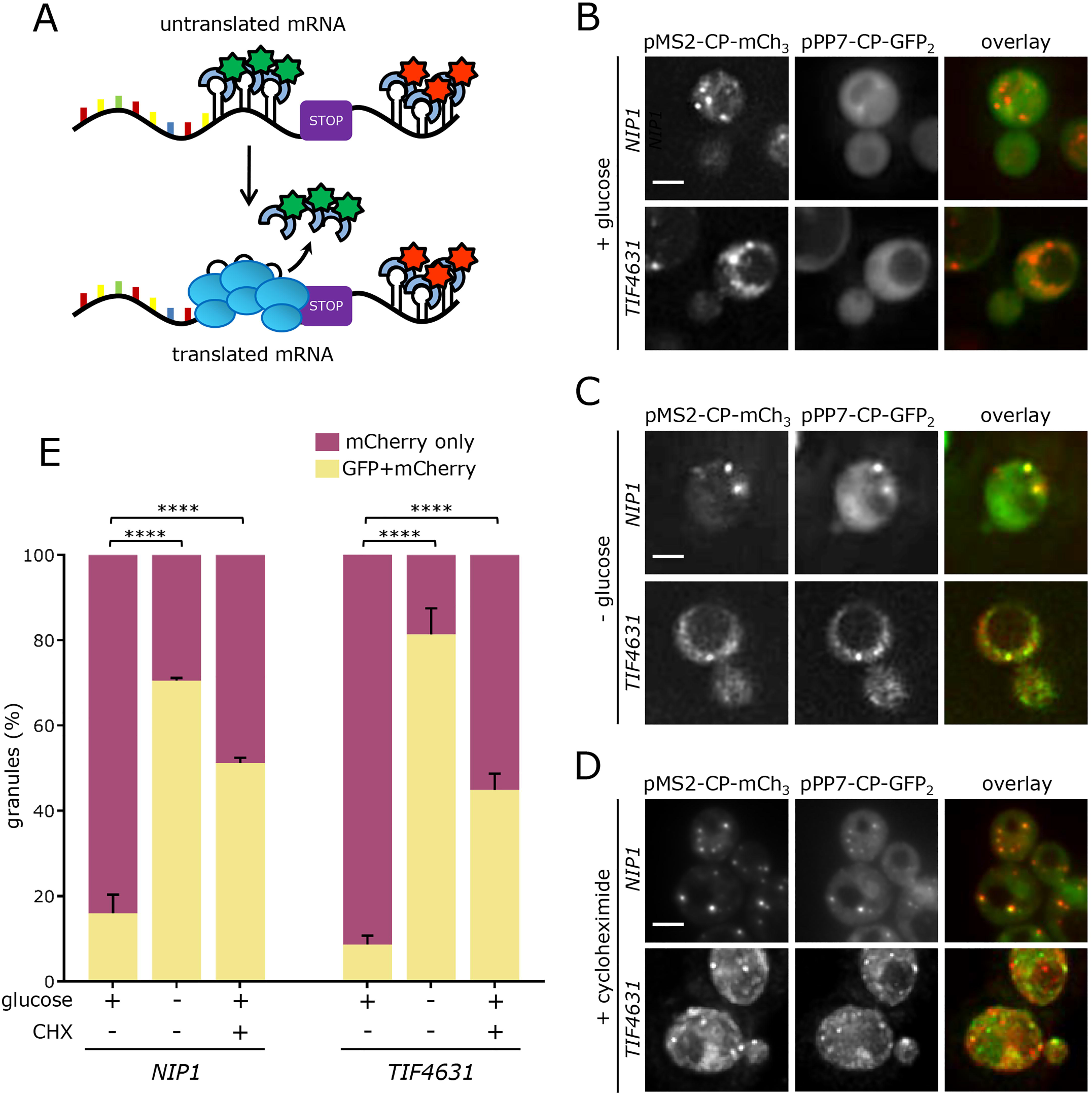
Granules house translationally active mRNAs. (A) Schematic representation of the TRICK strategy. (B) Z-stacked images showing *NIP1-TRICK* and *TIF4631-TRICK* in rich media, (C) following 10 minutes of glucose starvation or (D) after 10 minutes 100 μg/ml cycloheximide. (E) Chart showing the percentage of granules simultaneously showing GFP and mCherry signal under each condition. **** p<0.0005. Error bars indicate +SD. Scale bars = 4 μm.

A homologous recombination strategy was used to precisely insert a *TRICK* tag into the genome on the *TIF4631* or *NIP1* mRNAs. As above, the PP7 coat protein fused to GFP and the MS2 coat protein fused to mCherry were co-expressed and fluorescence was assessed. Under active growth conditions, mRNA granules can be observed for both the *NIP1* and *TIF4631* tagged mRNAs in the red but not the green fluorescent channel (Fig. 5B and 5F). This suggests that the MS2-mCherry fusion protein is bound to the mRNAs but that the PP7-GFP fusion is not bound (Fig. 5A). In contrast, after as little as 10 minutes glucose depletion, which leads to an almost total inhibition of translation initiation (Ashe et al., 2000), both fusion proteins are evident in granules (Fig. 5C and 5E). Similarly, cycloheximide treatment, which prevents ribosome translocation, also leads to an increase in the proportion of granules carrying both fluorescent protein fusions (Fig. 5D). This result mirrors what has been seen using the TRICK system in mammalian cells (Halstead et al., 2015). It seems likely that the cycloheximide causes decreased ribosomal transit without completely clearing ribosomes from the PP7 stem loop region. Therefore the level of PP7-GFP fusion protein binding induced by cycloheximide is lower than the level induced by glucose starvation, where ribosomal run-off is particularly extensive relative to most other stress conditions (Holmes et al., 2004). Overall, these data are highly suggestive that in live cells the translation factor mRNA granules are associated with active translation. This is analogous to our recent studies on mRNA granules housing two glycolytic mRNAs, where we found active protein synthesis was occurring possibly as a means to co-regulate protein production (Lui et al., 2014).

In sum, the data presented above suggest that translation factor mRNAs can be translated in granules and furthermore that this translation is a prerequisite for their localization. This localized translation likely occurs in a fluid phase-separated environment, such as has been described in the nucleolus, nuclear pore and p-granules (Brangwynne et al., 2009; Brangwynne et al., 2011; Frey et al., 2006).

### The Translation Factor mRNA granules are specifically inherited in a She2p/She3p dependent manner by the daughter cell

Whilst studying the localization of the mRNAs described above, it became clear that the granules harboring translation factor mRNAs were not evenly inherited during the yeast cell cycle suggesting that the location of protein production might provide the rationale for the mRNA localization. More specifically, mRNA granules harbouring the *NIP1* mRNA were observed to preferentially relocate into the developing daughter cell during cell division (Fig. 6A). Indeed, across hundreds of cell division events, preferential daughter cell re-localization of *NIP1* mRNA granules was observed in over 70% of cases (Fig. 6C). Equally, from the fixed cell *NIP1* smFISH studies roughly 55% of large multi-mRNA granules are found in developing buds, whereas the smaller single molecule mRNA foci are significantly less likely to be found at this location (Fig S7A)

**Figure 6.**
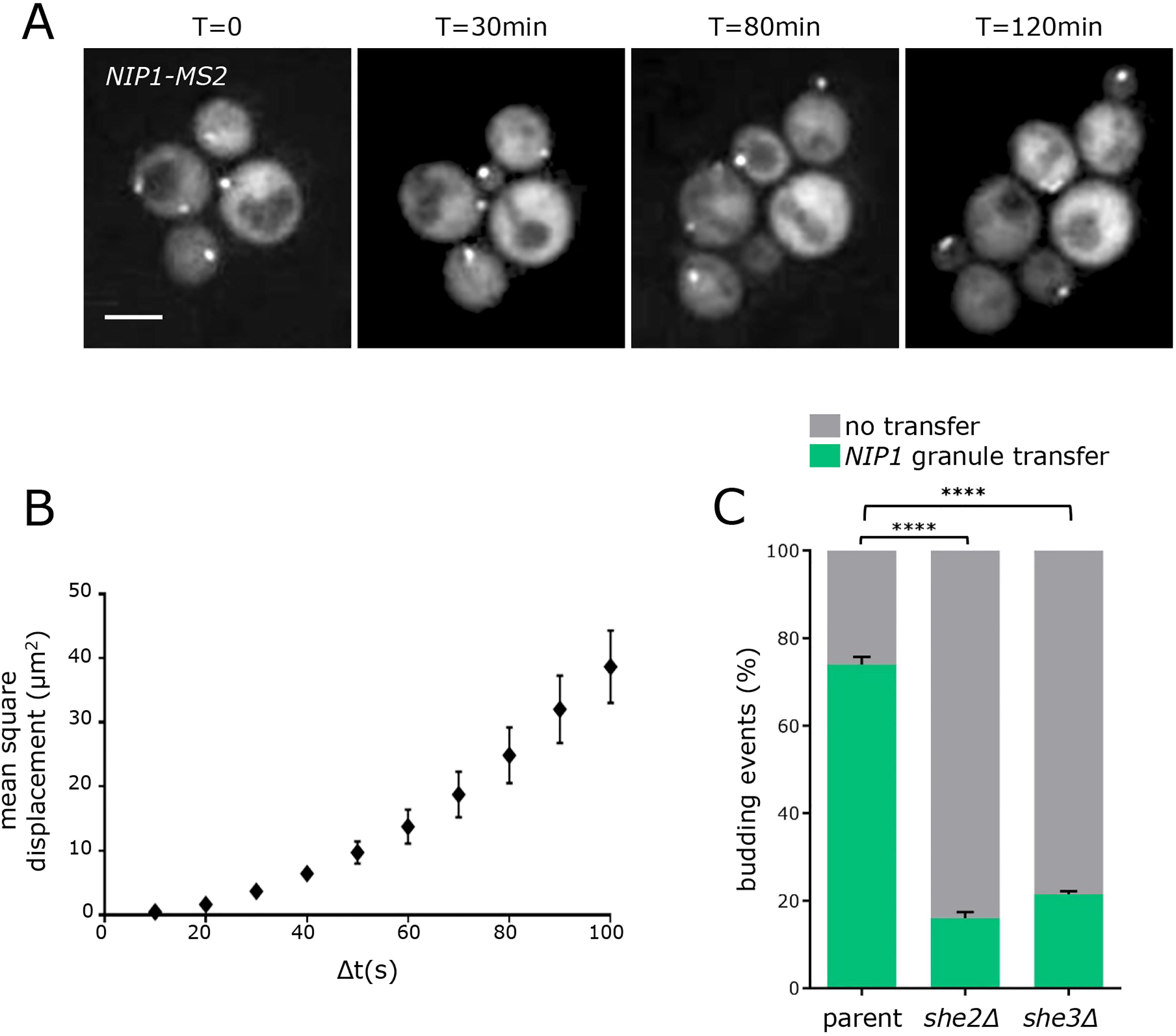
Granules move to the daughter cell upon division. (A) Z-stacked images of a *NIP1-MS2* strain bearing MS2-CP-GFP_3_ on a plasmid imaged over a period of two hours while under the constant flow of media. Scale bar = 4 μm. (B) Mean squared displacement analysis of *NIP1-MS2* granules, where the distance moved over increasing times is evaluated. Error bars indicate ± SD. (C) Chart showing the percentage of budding events in which transfer of a *NIP1-MS2* granule to the daughter cell occurs in wild type, *she2Δ* and *she3Δ* strains, calculated over three biological repeats. ****p<0.0001. Error bars indicate +SD.

A well-established method for evaluating whether particle movement is occurring via simple diffusion or in an activated manner is the mean squared displacement (MSD) plot (Qian, 2000). Time-lapse microscopy was therefore used to characterize the movement of the *NIP1* mRNA granules in cells by collecting images at 10 second intervals over a 2-minute period. The movement of single granules was tracked and used to generate MSD plots. In this common analysis (Qian, 2000), the average change in position of a molecule or body, known as the mean square displacement, is plotted over varying time intervals (Δt). The resulting curve provides information about the nature of the movement of a body or molecule within cells. Simple Brownian diffusion results in MSD values increasing linearly with Δt (Platani et al., 2002). Such a relationship was not observed for the plots generated from granules containing *NIP1* mRNA: instead, a distinct curve was evident (Fig. 6B). Similar curves have been associated with a combination of two or more types of movement (Platani et al., 2002) (Taylor et al., 2010). For instance, one possible explanation for this curve is that the granules oscillate between movable and non-movable phases possibly by being bound to transport machinery and a tether, respectively.

The yeast *ASH1* mRNA is well characterized as associated in tethered and movable states (Gonsalvez et al., 2004). It localizes specifically to the daughter cell as part of a translationally repressed RNP granule where it is tethered and translated. The machinery involved in the movement of this mRNA is particularly well characterised (Singer-Kruger and Jansen, 2014). For instance, She2p is an RNA-binding protein that specifically interacts with well-defined structures within the *ASH1* mRNA, and She3p is an intermolecular adaptor connecting She2p to the Myosin Myo4p, which moves along actin cables running from the mother cell to the daughter cell. In order to evaluate whether the same machinery is involved in the transit of translation factor mRNA granules, the *SHE2* and *SHE3* genes were deleted in strains carrying the *MS2*-tagged *NIP1* mRNA. Deletion of either gene led to the same effect, which was to dramatically reduce the level of mRNA granule transfer to daughter cells (Fig. 6C). Even though the machinery is the same as that involved in *ASH1* mRNA granules, the granules themselves do not co-localize (Fig. S7B). This observation is consistent with the difference in translational activity of mRNAs housed in these granules, with *ASH1* mRNA being repressed to prevent inappropriate expression in the mother cell during transit, whereas no such repression is evident for the translation factor mRNA granules.

The She2p/She3p machinery has also been implicated in the movement of mRNAs associated with the endoplasmic reticulum (Schmid et al., 2006). Therefore, it is possible that the translation factor mRNAs are also transported in association with the ER. If this were the case, the mRNA granules described above should at least partially overlap with ER localization. However, no such co-localization of ER and the *NIP1* mRNA granules was discernible (Fig. S7C). Equally, in previous datasets (Jan et al., 2014), translation factor mRNAs were not identified as enriched with ER (Fig. S7D). Similarly, *NIP1* mRNA granules did not appear to co-localize with mitochondria (Fig. S7E). However, it is still formally possible that the mRNAs are transported in a She2p-dependent manner while very transiently associated with an organelle such as the ER.

Overall, these data support a view that a She2p/She3p-dependent form of mRNA transit is employed in order that the translation factor mRNAs can be preferentially inherited by the daughter cell.

### The switch to filamentous growth is also associated with mRNA granule localization to the developing filamentous daughter cell

Given that a daughter cell will produce its own translation factor mRNAs and the maternal translated protein synthesis machinery is presumably free to diffuse within the cytosol of the mother or the developing daughter cell, it seems highly unlikely that there is an *absolute* requirement for polarization of translation factor mRNAs into the daughter cell. So why has such a mechanism evolved and what is the cellular benefit? Energetic considerations suggest that localizing mRNA rather than protein offers a significant advantage. Each mRNA molecule has been estimated to encode between 10^2^ and 10^6^ protein molecules, with average estimates between 1000-6000 protein molecules per mRNA(Futcher et al., 1999; Ghaemmaghami et al., 2003; Lawless et al., 2016; Lu et al., 2007). Clearly, robustly translated mRNAs will generate higher numbers of protein molecules, and in this case localizing an mRNA versus the several 1000 protein molecules it generates offers the cell significant energetic economies. However, in order that this energetic saving is realized, the protein synthetic machinery would also need to be localized to allow translation of the localized mRNAs. Furthermore, the polarization of mRNAs across cells might also relate to potential differing mRNA requirements of the daughter cell relative to the mother. Such a situation might be exacerbated when yeast respond to stress by inducing a different growth program, for example the switch from vegetative to filamentous growth.

Many laboratory strains have lost a capability that is evident in feral yeast strains to undergo filamentous growth patterns in response to different stress conditions (Liu et al., 1996; Lorenz et al., 2000). However, the Σ1278b strain can undergo filamentous growth in response to a range of nutritional stresses including nitrogen limitation, fusel alcohol addition and glucose depletion (Cullen and Sprague, 2012). Intriguingly, a *she2Δ* mutant in the Σ1278b strain is deficient in the switch from vegetative to filamentous growth and hence fails to undergo this form of polarization (Fig. 7A). It is entirely possible that a deficiency in the localization of translation factor mRNAs contributes to this phenotype.

**Figure 7.**
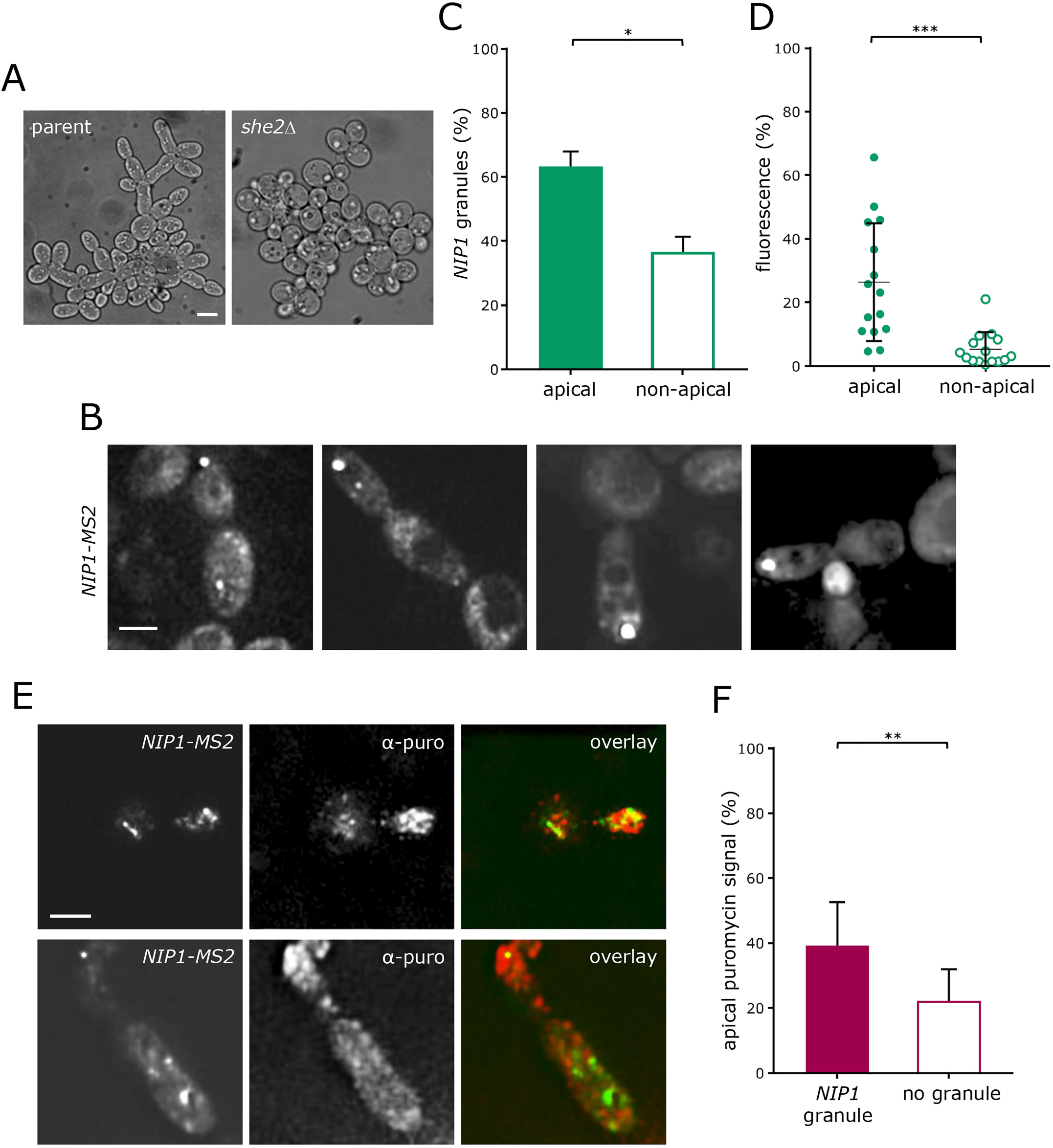
Granules localizing to the growing ends of cells growing as pseudohyphae. (A) Images of a *she2Δ* strain relative to the Σ1278b parent after growth in 1% butanol for 24 h (B) Z-stacked images of a Σ1278b strain showing filamentous phenotype after being grown in 1% butanol for 24 h. The strain expresses *NIP1-MS2* and MS2-CP-GFP_3_ from plasmids. (C) Chart showing the percentage of *NIP1-MS2* granules found in the apical quarter of an elongated cell, calculated over three biological repeats. *p<0.05. Error bars indicate +SD. (D) Chart showing the percentage of total cell fluorescence found in *NIP1* granules localising to the apical quarter compared to granules localising elsewhere. ***p<0.0005. Error bars indicate ±SD. (E) Z-stacked images of an immunofluorescence experiment performed on a 11278b strain bearing plasmid encoded *NIP1-MS2* and MS2-CP-GFP_3_ after treatment with puromycin. A primary antibody against puromycin and a Texas-red secondary antibody were used for immunofluorescence. (F) Graph showing the percentage of fluorescence intensity observed in the apical quarter of elongated cells that would either show (+) or not show (-) localisation of a *NIP1* granule to the apical quarter. **p<0.01. Error bars indicate +SD. Scale bars = 4 μm.

To explore the localization of mRNA during the switch to filamentous growth, *NIP1* mRNA granules were followed in Δ1278b strains treated with butanol to induce filamentation (Lorenz et al., 2000), and the granules were not only observed to preferentially localize to daughter cells but also to the most apical region of the daughter cell (Fig. 7B and C). Moreover, the granules found at this position showed on average a higher percentage of total cell fluorescence than granules found elsewhere in the cell (Fig. 7D) suggesting that a greater proportion of the mRNA localized to this region.

During filamentous growth, following commitment to a new cell cycle, yeast cells continue to grow apically from the growing tip instead of switching to isotropic growth, thus acquiring a characteristic elongated shape (Styles et al., 2013). It seems reasonable that continued apical growth might require a more intense rate of protein production at this site. Indeed, across a range of filamentous fungi, ribosomes or rough endoplasmic reticulum can be observed in extreme apical regions (Roberson et al., 2010). For instance, a subtending mass of ribosomes has been observed in the spitzenkörper of *Fusarium acuminatu* (Howard, 1981).

In order to assess whether more robust protein synthetic activity is observable near the apical tip of S. *cerevisiae* pseudohyphae, a yeast-adapted ribo-puromycilation assay (David et al., 2011; Lui et al., 2014) was performed on filamentous *S. cerevisiae* cells. In this assay, the addition of cycloheximide prevents polysomes runoff so that the translation machinery is locked on the transcript, while puromycin is added to the nascent polypeptide (David et al., 2012). Subsequent immunofluorescence for puromycin allows imaging of sites of global translation, while the GFP signal from the MS2-tagged mRNA is maintained throughout the procedure. This enables the simultaneous visualization of sites of protein production and *NIP1* mRNA granules. In this analysis, clouds of high puromycin signal were observed to surround prominent mRNA granules (Fig. 7E). It is important at this point to highlight the earlier result, that each granule likely contains a mixture of mRNAs. It is therefore reasonable to assume, when analyzing the localization of *NIP1,* that a number of other translation factor mRNAs might be present in the same location. Interestingly, the percentage of total puromycin signal in the apical quarter of the pseudohyphal cells was measured to be higher in cells carrying a *NIP1* mRNA granule in the same area than in cells showing a granule in other parts of the cell (Fig. 7F). These data are in accordance with the hypothesis that higher protein production rates are associated with the localization of translation factors to RNA granules.

## Discussion

In this study, we have identified and characterized a previously unanticipated localization for specific mRNAs that encode factors from the translation pathway. These mRNAs require translation for localization to granules and the granules themselves appear to represent sites of active translation. Single molecule studies show that approximately half of the molecules for each translation factor mRNA are present in the large multi-mRNA granules. These large mRNA granules localize specifically to the yeast daughter cell in a mechanism involving the She2p RNA-binding protein and the She3p-Myo4p binding protein. Furthermore, in polarized yeast cells undergoing filamentous growth, the translation factor mRNA granules localize to the apical region of the elongated daughter cell and this correlates with a region of high protein synthetic activity.

In previous work, we have used an MS2-tagging system and fluorescent *in situ* hybridization (FISH) to show that the transcript encoding elF4A *(TIF1)* was localized to granules in exponentially growing cells (Lui et al., 2014). Here, we again used the MS2-tagging system to show that mRNAs for various other factors involved in translation initiation, elongation and termination are localized to granules. Recent reports have highlighted that caution needs to be applied when interpreting live cell mRNA localization data using MS2-tethering approaches, as it is possible the *MS2* stem loops stabilize mRNA fragments and impact upon RNA processing (Garcia and Parker, 2015; Garcia and Parker, 2016; Haimovich et al., 2016; Heinrich et al., 2017). However, it has also been suggested that such phenomena are limited to a subset of transcripts and that such effects are more readily associated with plasmid-based expression systems (Haimovich et al., 2016). From our qRT-PCR data, it appears that the abundance of many of the transcripts analysed is not affected by insertion of the m-TAG, while for others the tagged version is significantly down-regulated relative to the endogenous version (Fig. SI). Given the concerns detailed above and the fact that stem loop insertion impacts upon the abundance of some of our tagged mRNAs, smFISH analysis was undertaken for endogenous untagged mRNAs. The data obtained accurately reproduce the localization patterns observed with the m-TAG system (Fig. 2). In addition, in previous studies the accumulation of MS2-derived mRNA fragments has been shown to coincide with Dcp2p containing foci or P-bodies (Haimovich et al., 2016). Under the active growth conditions used in our study, P-bodies are absent: therefore, the RNA granules do not colocalize with Dcp2p or P-bodies. These data agree with experiments where the insertion of poly(G) stem loops was necessary to observe the accumulation of mRNA 3’ fragments containing *MS2* stem loops under active growth conditions (Sheth and Parker, 2003). Under conditions that induce P-bodies, such as glucose depletion, co-localization of RNA granules with P-bodies can be observed (Lui et al., 2014; Simpson et al., 2014). Similar observations were made here for the translation factor mRNA granules suggesting an involvement of these mRNA granules in P-body formation where the mRNAs may get degraded as a consequence. Further evidence supporting the validity of the mRNA localization observed in this study stems from the fact that a variety of different transcripts exhibit different patterns of localization even though they all harbor the same *MS2* cassette. Some transcripts are not present in granules, some are present in 20 granules per cell and translation factor mRNAs are mostly present in less than 5 granules per cell. Furthermore, if the *MS2-* and *PP7*-tagging systems in dual-tagged strains were simply detecting mRNA fragments accumulating at sites of degradation these fragments should all co-localize. However, the data presented here show that the *MS2-* and *PP7*-tagged mRNAs overlap with one another to varying degrees: some overlap completely, some overlap partially, and some do not overlap at all. A final argument supporting the legitimacy of the mRNA localization data presented here comes from the ‘TRICK’ experiments performed for two transcripts. These data imply that the mRNAs in the granules are being translated, suggesting that the mRNAs are present in their full form. Therefore, while MS2-tethering strategies can impact upon various aspects of an mRNA’s fate, the approach does allow the investigation of RNA localization in live cells and permits an exploration of the altered localization under changing conditions. Fluorescent *in situ* hybridization approaches allow an investigation of the endogenous mRNA, but suffer from a need to fix cells; even if cellular fixation and permeabilization treatments don’t lead to alterations in mRNA pattern, FISH approaches do not allow the dynamics of mRNA localization to be studied in living cells.

Similarly to the granules housing two glycolytic mRNAs (Lui et al., 2014), the granules carrying translation factors described in this study appear to represent sites of active translation. Furthermore, the capacity of Pab1p to interact with poly(A) tails as well as the translation status of the mRNA seem fundamental for mRNA admittance into these granules. These data are suggestive of a scenario in which translation, or at least the potential for the mRNA to engage in translation, determines the capacity to enter the granule. Given that Pab1p interacts with the polyadenylation machinery, binds mRNA poly(A) tails in the nucleus and is likely exported with these transcripts (Brune et al., 2005; Dunn et al., 2005; Minvielle-Sebastia et al., 1997), it is possible that certain mRNPs are primed for entry into granules at this early stage. This could potentially offer an explanation as to why, for glycolytic mRNAs and the translation elongation factors mRNAs *TEF1* and *YEF3,* the levels of co-localization within granules mirrors similarities in transcription patterns. Indeed, increasing evidence points to inherent connections between the nuclear history of a transcript and its cytosolic fate (Bregman et al., 2011; Gunkel et al., 1995; Trcek et al., 2011; Zid and O’Shea, 2014).

Interestingly, the translationally active state of mRNAs within the granules is very rapidly reversed upon glucose starvation: a condition known to induce the rapid formation of P-bodies after translation inhibition. In such conditions, the degree of overlap among different mRNAs in the granules increases strikingly, in accordance with the observation that distinct granules coalesce during the formation of P-bodies (Lui et al., 2014). Considering that yeast P-bodies have recently been described as liquid-like droplets (Kroschwald et al., 2015) and that the granules described in this work seem to be similarly sensitive to hexanediol treatment, it is not difficult to imagine how the transition from translation granules to P-bodies could occur; especially given that the rapid assembly and exchange of components are facilitated within bodies with liquid-like properties (Kroschwald et al., 2015). One intriguing explanation as to how the granules might coalesce when forming P-bodies is that a glucose starvation-induced ‘contraction’ of the cytosol (Joyner et al., 2016) might induce fusion of the granules by simple molecular crowding effects, or as a consequence of an altered phase separation between the granules and the cytosol.

What emerges from these observations is a scenario in which certain mRNAs seem to exist in RNP granules where they can either undergo translation or decay, depending on the requirements of the cell. The presence of RNA-containing granules associated with degradation or mRNA localization is very widely reported, where such granules are generally associated with translational repression, while the potential for specialized translation foci is less widely acknowledged. One obvious advantage to the co-localization of mRNAs is the potential for co-translational folding of protein-protein complexes (Shiber et al., 2018). Indeed, many of the translation initiation factors are present as complex multi-subunit factors. For example, we have investigated the localization of components of elF2B, elF2 and elF3, and one obvious possibility is that these complexes are constructed co-translationally.

In recent years, there has been an increased appreciation of the relationship between mRNA co-localisation and protein complex formation. For example in yeast, from a study of 12 multi-subunit protein complexes, 9 were shown to form co-translationally (Shiber et al., 2018). Likewise in human cells, the dynein heavy chain mRNA co-localizes at sites of translation, possibly as a way to facilitate assembly of the mature protein complex (Pichon et al., 2016). Similarly, mRNAs for many of the components of the Arp2/ Arp3 complex are localized and co-translated at the leading edge of fibroblasts possibly to aid in protein complex formation (Mingle et al., 2005; Willett et al., 2013). Equally, the peripherin mRNA has previously been proposed to localize to specialized factories that couple the localization and translation of the transcript with the assembly of peripherin intermediate filaments, in a process termed dynamic co-translation (Chang et al., 2006).

A key feature of all these examples is the necessity for translation of classes of mRNAs in a distinct region of the cell and hence the presence of the translation machinery at this locale. The concentration of the translation machinery in certain areas of eukaryotic cells has previously been associated with asymmetric growth: in migrating fibroblasts, translation factors were found to preferentially localize to the lamellipodia, where the rates of protein production are higher (Willett *et al.,* 2010; Willett *et al.,* 2011). Furthermore, local translation was identified as a key regulator of cellular protrusions in migrating mesenchymal cells (Mardakheh et al., 2015).

In this study, we show that translation factor mRNA granules are transported to the daughter cell in a She2p/She3p/Myo4p machinery-dependent manner. The specific localization of translation factor mRNAs to the daughter cell provides a compelling rationale for the RNA granules, in that they might provide the daughter cell with a ‘start-up’ pack concentrating protein synthetic activity to facilitate daughter cell development. Given that approximately half the molecules of each individual mRNA are present in such granules, a mother cell is essentially donating half of its mRNA to the developing daughter cell. Such an idea has parallels with maternal inheritance of mRNA by the oocyte in multicellular organisms such as Xenopus and Drosophila (Lee et al., 2014). We propose that the granules represent specialized factories for the translation machinery, which are specifically inherited by the daughter cell. As such, protein synthetic activity would be converged to an area of the cell where it is particularly required.

## Materials and Methods

### Strains and plasmids

The S. *cerevisiae* strains used in this study are listed in Table I. *MS2* and *PP7* stem loops were PCR amplified from the pLOXHIS5MS2L and pDZ416 plasmids respectively using primers directed to the 3’ UTR of the relevant genes. After transformation and selection, accurate homologous recombination of the resulting cassette was verified using PCR strategies and the selection marker was subsequently excised using Cre recombinase. pMS2-CP-GFP_3_, pMS2-CP-mCherry_3_ or pMet25MCP-2yEGFP (pDZ276) plasmids were then transformed into the strains to enable detection of *MS2* and *PP7*-tagged mRNAs The *MS2* and *PP7* tagging reagents were gifts from Jeff Gerst and Robert Singer (Addgene #31864 & #35194) (Haim-Vilmovsky and Gerst, 2009; Hocine et al., 2013). Dual *MS2-* and *PP7*-tagged strains were obtained by mating of appropriate haploid strains, followed by sporulation and tetrad dissection. TRICK strains were generated using a similar approach to above, but using a DNA template developed for TRICK in yeast. Briefly, a 12x*PP7* 24*xMS2* synthesized fragment (Halstead et al., 2015) was subcloned into the pFA6a-kanMX6 vector and specific targeting primers were used to isolate the TRICK region with the marker gene such that integration into the *NIP1* and *TIF4631* genes was achieved. For *she2Δ* and *she3Δ* strains, the ORFs were replaced by the nourseothricin resistance gene, *(natNT2)* amplified from the pZC2 vector (Carter and Delneri, 2010). A *PAB1* shuffle strain was generated in the yMK2254 *NIP-MS2* strain by first transforming a *PAB1 URA3* plasmid then deleting the *PAB1* gene with a *LEU2* cassette. *PAB1* mutant strains were generated by transformation of *PAB1-ΔRRM2 TRP1* and *PAB1-Y83V,F170V TRP1* plasmids (Kessler and Sachs, 1998) into the shuffle strain followed by expulsion of the *PAB1 URA3* plasmid. For generation of the yEPIacl95-*NIP1* plasmid, *MS2-*tagged *NIP1* was amplified from the yeast strain yMK2254 and cloned into yEPIacl95 (Gietz and Sugino, 1988). A stem loop sequence (Vattem and Wek, 2004) was inserted into this plasmid using a PCR-based approach, where the stem loop was introduced on primers that directed amplification of the entire plasmid which was subsequently verified by DNA sequencing.

**Table I.** Yeast strains used in this study.

| Strain | Genotype | Source |
| --- | --- | --- |
| yMK7 | *MATa leu2Δ::hisG his3Δ::hisG trp1Δ::hisG ura3-52* | JCY100 (J. Thorner) |
| yMK467 | *MATα ADE2 his3-11,15 leu2-3,112 trp1-1 ura3-1* | Ashe strain collection |
| yMK807 | *MATa ADE2 his3-11,15 leu2-3,112 trp1-1 ura3-1* | Ashe strain collection |
| yMK1585 | yMK467 *TIF1-MS2L p[MS2-GFP<sub>3</sub> HIS3]* | Ashe strain collection |
| yMK1741 | yMK467 *p[MS2-GFP<sub>3</sub> HIS3]* | Ashe strain collection |
| yMK1833 | yMK466 *CDC33-RFP::NAT DCP2-CFP::TRP1 NPC2-MS2L p[MS2-GFP<sub>3</sub> HIS3]* | Ashe strain collection |
| yMK2124 | yMK467 *SUP35-MS2L p[MS2-GFP<sub>3</sub> HIS3]* | This study |
| yMK2134 | yMK467 *GCD6-MS2L p[MS2-GFP<sub>3</sub> HIS3]* | This study |
| yMK2136 | yMK467 *GCD7-MS2L p[MS2-GFP<sub>3</sub> HIS3]* | This study |
| yMK2218 | yMK466 *TIF1-PP7L p[PP7-GFP<sub>2</sub> URA3]* | This study |
| yMK2249 | yMK467 *CDC33-MS2L p[MS2-GFP<sub>3</sub> HIS3]* | This study |
| yMK2251 | yMK466 *ENO2-PP7L p[PP7-GFP<sub>2</sub> URA3]* | This study |
| yMK2254 | yMK467 *NIP1-MS2L p[MS2-GFP<sub>3</sub> HIS3]* | This study |
| yMK2272 | yMK467 *TIF4632-MS2L p[MS2-GFP<sub>3</sub> HIS3]* | This study |
| yMK2362 | yMK467 *EFB1-MS2L p[MS2-GFP<sub>3</sub> HIS3]* | This study |
| yMK2363 | yMK467 *YEF3-MS2L p[MS2-GFP<sub>3</sub> HIS3]* | This study |
| yMK2364 | yMK467 *TIF4631-MS2L p[MS2-GFP<sub>3</sub> HIS3]* | This study |
| yMK2365 | yMK466 *NIP1-MS2L TIF1-PP7L p[MS2-mCh<sub>3</sub> HIS3] p[PP7-GFP<sub>2</sub> URA3]* | This study |
| yMK2369 | yMK467 *she2::NAT NIP1-MS2L p[MS2-GFP<sub>3</sub> HIS3]* | This study |
| yMK2370 | yMK467 *she3::NAT NIP1-MS2L p[MS2-GFP<sub>3</sub> HIS3]* | This study |
| yMK2372 | yMK466 *TIF4631-MS2L TIF1-PP7L p[MS2-mCh<sub>3</sub> HIS3] p[PP7-GFP<sub>2</sub> URA3]* | This study |
| yMK2373 | yMK466 *EFB1-MS2L TIF1-PP7L p[MS2-mCh<sub>3</sub> HIS3] p[PP7-GFP<sub>2</sub> URA3]* | This study |
| yMK2519 | yMK467 *TEF1-MS2L p[MS2-GFP<sub>3</sub> HIS3]* | This study |
| yMK2542 | yMK466 *TIF4632-PP7L p[PP7-GFP<sub>2</sub> URA3]* | This study |
| yMK2564 | yMK7 *p[NIP1-MS2L] p[MS2-GFP<sub>3</sub> HIS3]* | This study |
| yMK2567 | yMK467 *pab1::LEU2 NIP1-MS2L p[MS2-GFP<sub>3</sub> HIS3] p[PAB1 TRP1]* | This study |
| yMK2614 | yMK467 *SUI2-MS2L p[MS2-GFP<sub>3</sub> HIS3]* | This study |
| yMK2616 | yMK467 *pab1::LEU2 NIP1-MS2L p[MS2-GFP<sub>3</sub> HIS3] p[PAB1-ΔRRM2 TRP1]* | This study |
| yMK2617 | yMK467 *pab1::LEU2 NIP1-MS2L p[MS2-GFP<sub>3</sub> HIS3] p[PAB1-Y83V,F170V TRP1]* | This study |
| yMK2672 | yMK467 *TEF1-MS2L ENO2-PP7L p[MS2-GFP<sub>3</sub> HIS3] p[PP7-GFP<sub>2</sub> URA3]* | This study |
| yMK2686 | yMK467 *TIF4631-TRICK-stop-MS2L p[MS2-mCh<sub>3</sub> HIS3] p[PP7-GFP<sub>2</sub> URA3]* | This study |
| yMK2687 | yMK467 *DCP2-CFP::TRP1 NIP1-MS2L TIF1-PP7L p[MS2-mCh<sub>3</sub> HIS3] p[PP7-GFP<sub>2</sub> URA3]* | This study |
| yMK2688 | yMK467 *NIP1-TRICK-stop-MS2L p[MS2-mCh<sub>3</sub> HIS3] p[PP7-GFP<sub>2</sub> URA3]* | This study |
| yMK2941 | yMK467 *p[NIP1-MS2 URA3] p[MS2-GFP<sub>3</sub> HIS3]* | This study |
| yMK2942 | yMK467 *p[sl-NIP1-MS2 URA3] p[MS2-GFP<sub>3</sub> HIS3]* | This study |
| yMK2949 | yMK467 *TIF4631-MS2L TIF4632-PP7L p[MS2-mCh<sub>3</sub> HIS3] p[PP7-GFP<sub>2</sub> URA3]* | This study |

### Yeast growth

Strains were grown at 30°C on Synthetic Complete medium with 2% glucose (SCD) with selection where necessary (Sherman, 1991). Cells were incubated for 30 min in SCD media lacking methionine to induce expression the CP-GFP/RFP fusions prior to imaging. For experiments requiring glucose starvation, exponentially growing cells were resuspended in media lacking glucose, then incubated for 10 min at 30°C before imaging. For induction of filamentous growth, the JCY100 strain (Σ1278b background) (Cook et al., 1997) was grown in SCD media containing 1% butanol for up to 24 h at 30°C prior to imaging.

### Fluorescent microscopy

Live cell microscopy was performed on a Delta Vision microscope (Applied Precision) equipped with a Coolsnap HQ camera (Photometrics), using a 100x/ 1.40 NA oil plan Apo objective. Imaging was performed for GFP (excitation-490/20 nm, emission-535/50 nm, exposure-200-400ms), mCherry (excitation-572/35 nm, emission-632/60 nm, exposure-400-800ms) and CFP (excitation-436/10 nm, emission-465/30, exposure-). Images were acquired using Softworx 1.1 software (Applied Precision) and processed using Image J software package (National Institute of Health, NIH). For routine live-cell imaging, exponential yeast were viewed on poly-L-lysine coated glass slides. For live cell imaging over longer periods of time, a microfluidic system (CellASIC) (Merck Millipore) was used, where exponential yeast were imaged every of 10 min for 2 h. For smiFISH, images of fixed samples were collected on a Leica TCS SP8 AOBS inverted gSTED microscope using a 100x/1.40 Plan APO objective and 1x confocal zoom. The confocal settings were as follows, pinhole 1 airy unit, scan speed 400Hz bidirectional, format 1984 x 1984. DAPI images were collected using a photon multiplying tube detector, with a blue diode 405nm laser (5%). Confocal images were collected using hybrid detectors with the following detection mirror settings; Alexa Fluor 488 410-483nm (5 to 50μs gating); Alexa Fluor 546 556-637nm (5 to 35μs gating); Alexa Fluor 647 657-765nm **(5-50μS** gating) using the 488nm (60%), 546nm (60%) and 646nm (60%) excitation laser lines, respectively. Images were collected sequentially in 200nm Z sections. Acquired images were subsequently deconvolved and background subtracted using Huygens Professional (Scientific Volume Imaging).

### smFISH and Immunofluorescence

For smFISH, gene specific 20nt antisense oligonucleotides were designed with a 5’ Flap sequence, to which fluorescently labelled oligonucleotides were annealed (Tsanov et al., 2016). 30-48 probes were designed per mRNA such that each probe had minimal potential for cross-hybridisation and between 40 and 65% GC content (probe sequences are available upon request). To generate the fluorescently labelled smFISH probes, 200pmoles of an equimolar mix of gene specific oligos was annealed with 250pmoles of the appropriate fluorescently labelled flap oligo (Integrated DNA Technologies), as described previously (Tsanov et al., 2016). To perform smFISH, strains were grown in SCD overnight to mid-log phase and fixed with 4% EM grade formaldehyde (Electron Microscopy Sciences 15714-S) for 45 minutes, at room temperature. After fixation, cells were washed with buffer B (1.2M Sorbitol, 100mM KHPO_4_, pH 7.5), then resuspended in spheroplasting buffer (1.2M Sorbitol, 100mM KHPO 4, 20 mM Ribonucleoside Vanadyl Complex (VRC), 0.2% β-mercaptoethanol, 1 mg/ml lyticase) and incubated at 37°C for 15 minutes before being permeabilized with 70% Ethanol. Subsequently, cells were hybridized with 20pmoles of the appropriate fluorescently labeled smFISH probes in hybridization buffer (10mg *E. coli* tRNA, 2mM VRC, 200 μg/ml BSA, 10% Dextran Sulfate, 10% Formamide, 2X SSC in nuclease free water). Cells were then washed in 10% Formamide, 2X SSC and adhered to 0.01% Poly-L-Lysine coated coverslips before mounting in ProLong™ diamond antifade mountantwith DAPI (Life Technologies).

For immunofluorescence, cells were grown to mid-log phase in media with 1M sorbitol, incubated for 1 h with 1 mg/ml lyticase, then incubated for 20 min with 1 mg/ml puromycin and 100 μg/ml cycloheximide. Cells were then fixed in 4% formaldehyde and loaded on poly-L-lysine coated coverslips. Coverslips were blocked for 30 min in 4% Bovine Serum Albumin then incubated overnight with a mouse anti-puromycin monoclonal antibody (Millipore) (1:1000 in 4% BSA). After a lx PBS wash, coverslips were incubated with an anti-mouse Texas Red-conjugated secondary antibody (Abeam) (1:200 in 4% BSA) for 2 h, then mounted and imaged.

### Quantification and statistics

For quantification of granule numbers per cell, 100 cells were counted for each strain across 3 biological repeats. For quantification of overlapping MS2 and PP7 signal in double-tagged strains or TRICK strains, 100-150 granules were considered for each strain over three biological repeats. For quantification of budding events and the inheritance of granules, all the budding events observable (approx. 30) over 3 different frames were considered for each strain over 3 biological repeats. For quantification of granules found in the apical quarter of filamentous cells, the length of the cell was calculated using ImageJ and granules found within a quarter of the length from the apical end were counted. Three biological repeats were considered, with at least 150 cells counted per repeat. For quantification of percentage of fluorescence, the intensity of fluorescence was measured using ImageJ for 15 cells. The corrected total fluorescent intensity for the whole cell and for the granules was measured to calculate the percentage of fluorescence in granules. GraphPad Prism 7 (GraphPad Software, Inc.) was used to produce the graphs and to calculate the standard error of the mean, indicated by error bars. Two-way ANOVA was performed using GraphPad Prism 7.

SmFISH micrographs were analyzed using FISHQuant (Mueller et al., 2013) and FindFoci (Herbert et al., 2014) to provide sub-pixel resolution of spot locale and spot enhancement via dual Gaussian filtering. The resulting output files were then processed using custom scripts in R to assess spot co-localisation, mRNA copies per spot and mRNA copies per cell. For spot co-localization analysis, each spot in one channel was paired with the closest spot in the opposite channel based on spot centroid distance in 3D space. Spots were deemed to co-localize if the 3D distance between them was less than the summed radius of the two spots. To assess the number of mRNAs in each spot, the cumulative fluorescent intensity of all spots was calculated and fit to a Gaussian curve, the peak of which corresponds to the intensity of a spot containing a single mRNA (Trcek et al., 2017). This value was used to normalize the cumulative intensity of each spot, thus determining the number of mRNAs per spot (Trcek et al., 2012). Subsequently, the mean number of mRNAs per cell was calculated using these values and cross-compared with values obtained from genomic studies using RNA-seq (Lahtvee et al., 2017; Lawless et al., 2016). For micrograph pseudo-colouring, foci were assigned a grayscale value corresponding to the number of predicted mRNAs within that spot, calculated as above. Subsequently, these grayscale intensities were ‘colored’ and visualised using a custom LUT.

### Mean square displacement

Strain yMK2254 was imaged at intervals of 10 s over a total time of 2 min. Granules were followed and the distance moved was measured using ImageJ software (NIH). The distances travelled by granule in 10 s intervals were used to calculate the Mean Square Displacement (MSD) using the equation, MSD (Δt) = [d(t) - d(t+ Δt)]^2^; where Δt = time interval between images, and d(t) = the position of the RNA granule at a given time t (Platani et al., 2002).

### Quantitative RT-PCR

To extract RNA, 50 ml mid-log phase yeast cultures were pelleted, resupended in 1 ml Trizol (thermofisher scientific) then 400μl acid washed beads (sigma) were added. Tubes were sequentially vortexed five times for 20 s with 1 min intervals. 150μl chloroform was added and the samples were mixed. The tubes were centrifuged in a microfuge for 15 min at 12000xg. The aqueous layer was collected and 350μl isopropanol was added. The resulting precipitate was collected via centrifugation in a microfuge for 15 min at 12,000xg and washed in 75% ethanol. The resulting pellet was resuspended in 20μl of nuclease-free H_2_0. Quantitative RT-PCR (qRT-PCR) was performed using 300 ng RNA with the CFx Connect Real-Time system with the iTaq Universal SYBR Green One Step Kit (Bio-Rad) according to manufacturer’s instructions. Primers were designed to amplify a 200n region just upstream of the STOP codon. Samples were run in triplicate and normalized to *ACT1* mRNA, and the fold change was calculated using 2^-ΔCt^ for each tested RNA.

## Acknowledgements

We thank G. Pereira, J. Gerst, J. Thorner, J. Chao and R. Singer for reagents. M.P. was supported by a Wellcome Trust PhD studentship (099732/Z/12/Z). C.B., E.L. and B.W. were supported by Wellcome Trust PhD studentships (210002/Z/17/Z, 210004/Z/17/Z and 109330//Z/15/Z). J.L. and G.F. were supported by a Biotechnology and Biological Sciences Research Council (BBSRC) project grant (BB/K005979/1), F.M-P. was supported by a CONICYT Becas Chile PhD studentship (ref. 72140307). The Bioimaging Facility microscopes used in this study were purchased with grants from BBSRC, Wellcome Trust and the University of Manchester Strategic Fund. Thanks go to Peter March and Steve Marsden for their help with the microscopy.

## Author contributions

All authors generated reagents, performed experiments and evaluated results. M. Pizzinga, C. Bates and M. Ashe conceived the study, proposed and designed experiments and wrote the manuscript. All authors contributed to the discussion and evaluation of the manuscript.

