## Supplementary figures and images for "Translation factor mRNA granules direct protein synthetic capacity to regions of polarized growth"

### Supplemental figure 2

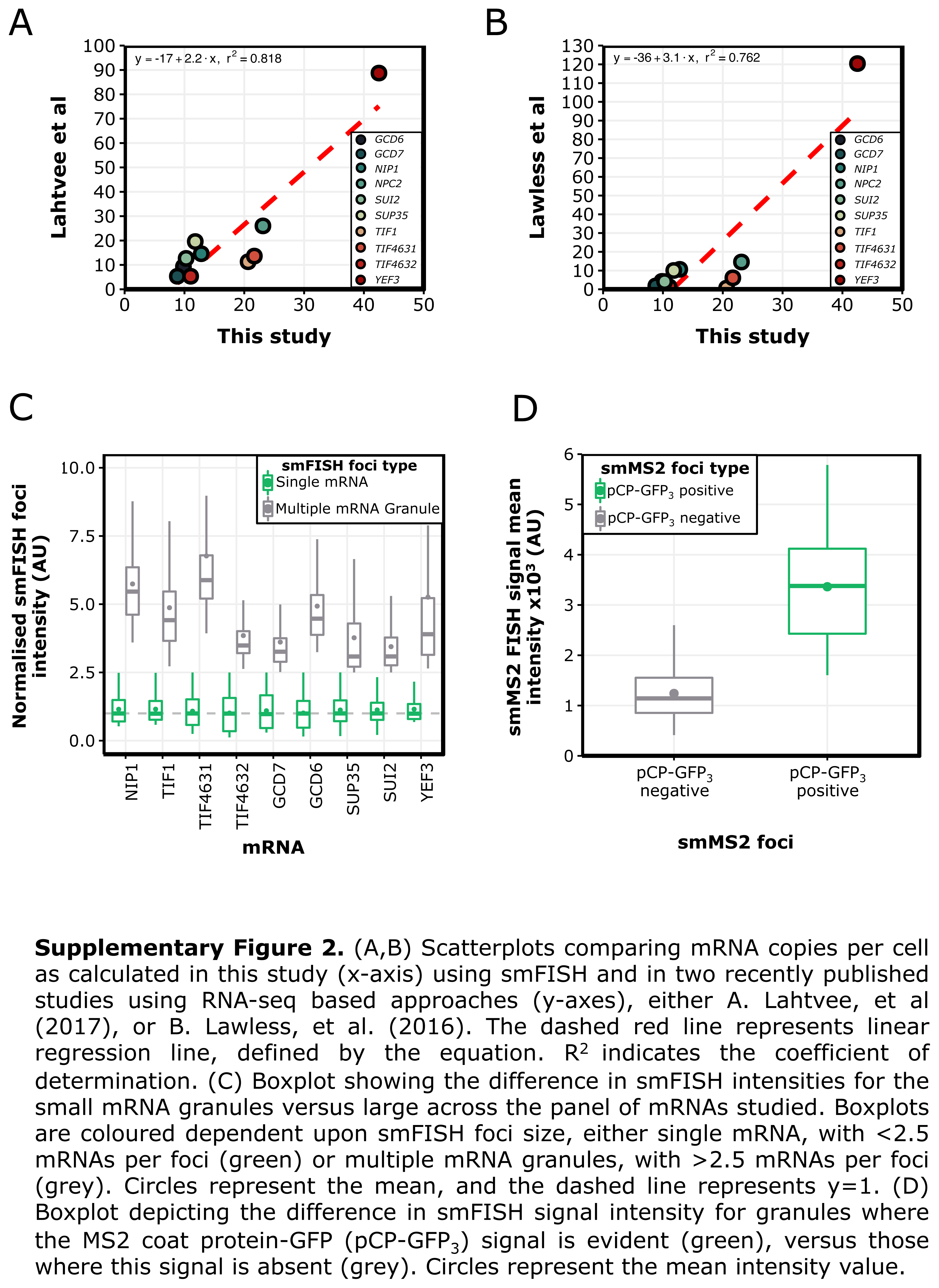

### Supplemental Figure 6

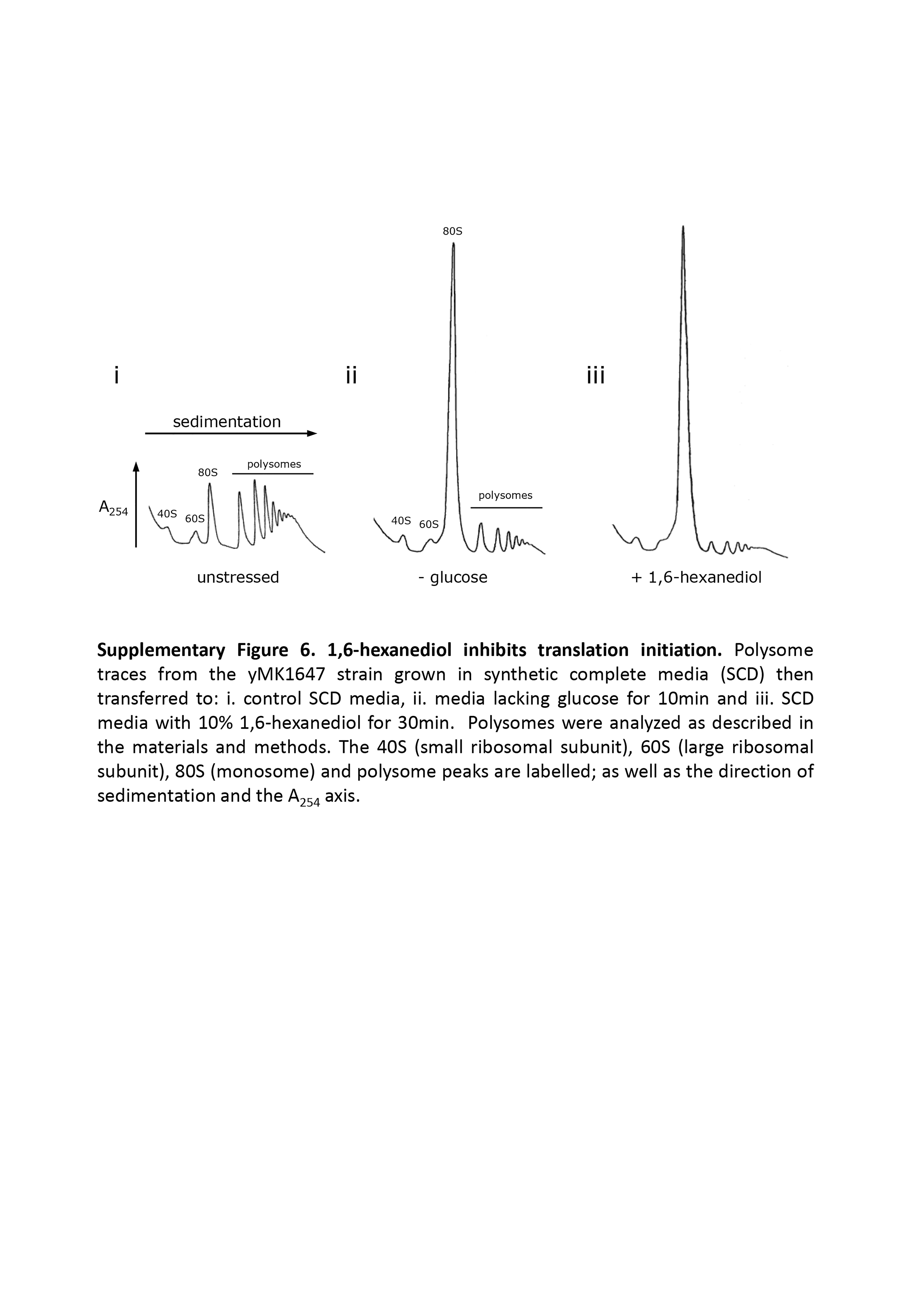

### Supplemental Figure 7

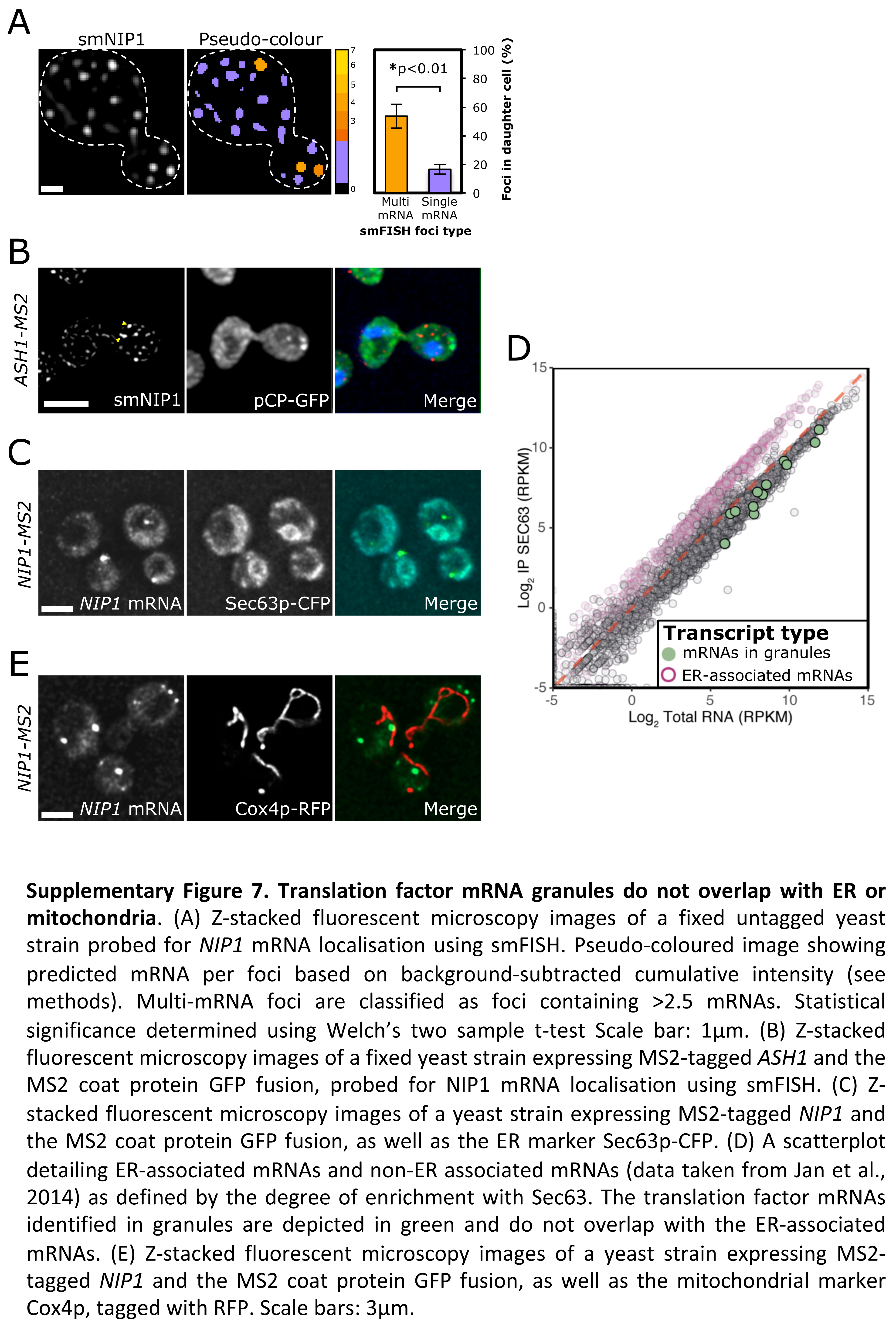
